# Queuosine glycosylation occurs after anticodon loop methylations and protects against stress-induced tRNA fragmentation

**DOI:** 10.64898/2026.06.28.735034

**Authors:** Ana C. de A. P. Schwarzer, Julia Dietzsch, Sabrina Stille, Nicolas Lemus-Diaz, Milán Erich, Eva Schöller, Katharina Zimmerningkat-Sievers, Achim Dickmanns, Ralf Ficner, Katherine E. Bohnsack, Claudia Höbartner, Markus T. Bohnsack

## Abstract

Queuosine, present at position 34 (Q_34_) of vertebrate tRNA^Asp^ and tRNA^Tyr^, is mannosylated and galactosylated by QTMAN and QTGAL, respectively. However, the interplay between these glycosylations and other anticodon loop modifications, and their impact on tRNA fragmentation, have remained unknown. Here, we define hierarchical modification pathways for the anticodon loop modifications of the two human glycoQ_34_-containing tRNAs. Our data reveal bidirectional interdependence of Q_34_ and m^5^C_38_ in tRNA^Asp^ with subsequent Q_34_ mannosylation occurring in an m^5^C_38_-independent manner. Anticodon loop modifications of tRNA^Tyr^ are installed independently, but sequentially in a pathway in which TRMT5-mediated m^1^G_37_ formation precedes queuosinylation and galactosylation. Q_34_ glycosylation enhances protein synthesis but alongside their canonical function in translation, tRNAs are precursors of tRNA-derived fragments that can regulate diverse aspects of gene expression. Our data reveal that Q_34_ mannosylation protects tRNA^Asp^ from stress-induced cleavage and modulates cleavage site selection, thus expanding the relevance of Q glycosylation beyond translation.

## INTRODUCTION

Cellular RNAs contain a plethora of modified nucleotides that contribute to their functions and regulation^1^. RNA modifications can be introduced post-transcriptionally through the addition of chemical groups, such as methylations, to various positions of the canonical nucleotides adenosine (A), uracil (U), cytidine (C) and guanosine (G)^2^. However, canonical nucleotides can also be converted into, or exchanged for, non-canonical nucleotides, with A-inosine editing^3^, isomerization of U to pseudouridine^4^, and the swapping of G for queuosine (Q)^5^ being prominent examples. Like the canonical nucleotides, non-canonical nucleotides in RNAs can also be substrates for the addition of chemical groups, wherein hypermodifications that have recently come into focus are the mannosylation and galactosylation of Q to form manQ and galQ, respectively^6^.

In eukaryotes, Q is incorporated at position 34 of tRNAs with “GUN” anticodons (aspartate (Asp), asparagine (Asn), histidine (His), and tyrosine (Tyr)) by the tRNA guanine transglycosylase (TGT), a dimeric enzyme composed of the QTRT1 and QTRT2 subunits^6–8^. Eukaryotes do not produce Q *de novo*, but rather take up the queuine (q) base from nutritional sources or from microbiota, and then Q is directly incorporated into the target tRNAs via transglycosylation^7,9^. In this reaction, a conserved Asp residue in TGT performs a nucleophilic attack that results in a temporary covalent bond with the ribose 34, excising the original G, which is released from the active site^9^. Upon binding of queuine, the base is activated by a second Asp residue in TGT, and a new glycosidic bond is formed with ribose 34 thus incorporating the non-canonical base^9^. The catalytic functions are performed by QTRT1, whereas the inactive QTRT2 subunit is required for interactions with the substrate, positioning the tRNA and assisting conformational rearrangements of the anticodon loop to reach the active site in QTRT1^9–11^. Some of the known recognition elements present in TGT substrates include the G_34_ and U_35_ residues, as well as the stem loop structure of the tRNA anticodon^9,11,12^. Following queuosinylation, Q_34_ of tRNA^Asp(GUC)^ and tRNA^Tyr(GUA)^ are targeted by the RNA glycosylases QTMAN and QTGAL, respectively^13^ (**Fig. 1A**). QTMAN transfers a mannose from GDP-mannose to the dihydroxycyclopentene ring of Q to form an a-glycosidic bond with the allylic hydroxy group, whereas QTGAL catalyzes an S_N_2-type reaction to form a b-glycoside in which the homoallylic hydroxygroup of Q obtains a galactose from UDP-galactose^13–15^. In tRNA^Tyr(GUA)^, Q can be incorporated and glycosylated prior to splicing, indicating that these are early tRNA biogenesis events^16^.

**Figure 1.**
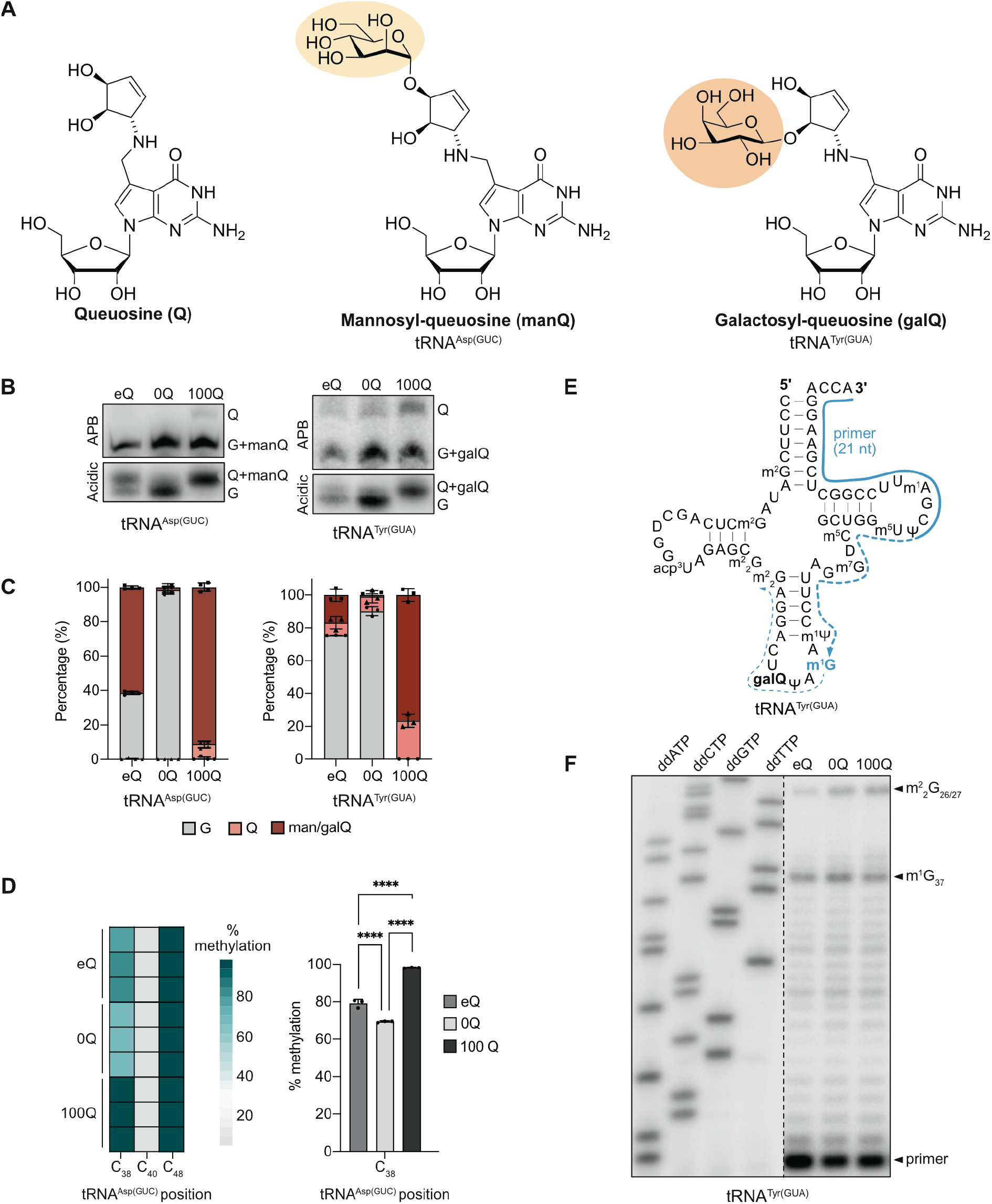
Levels of Q_34_ and man/galQ_34_ in tRNA^Asp(GUC)^ and tRNA^Tyr(GUA)^ and influence of queuosinylation on m^5^C_38_/m^1^G_37_. **(A)** Chemical structures of Q, manQ and galQ with glycosylations highlighted in yellow/orange. **(B)** Total RNAs from HEK293 cells grown in the presence of FBS (endogenous (e)Q), horse serum (0Q) or in horse serum supplemented with q (100Q) were separated by APB (5%) or acidic gel electrophoresis and tRNA^Asp(GUC)^/tRNA^Tyr(GUA)^ were detected by northern blotting. Representative image of n=4 independent experiments is shown. **(C)** Quantification of the levels of G_34_, Q_34_ and man/galQ_34_ in samples as described in (B). Error bars represent mean ± standard deviation. **(D)** RNAs from eQ, 0Q and 100Q cells were subjected to bisulfite sequencing to monitor m^5^C_38_ levels in tRNA^Asp(GUC)^. Results from n=3 independent experiments are shown as a heatmap (left) and a bar plot (right; error bars represent mean ± standard deviation). **(E)** Scheme of primer extension assay to detect m^1^G_37_ and m^2^_2_G_26/27_ in tRNA^Tyr(GUA)^. **(F)** Primer extension assay monitoring m^1^G_37_ levels in tRNA^Tyr(GUA)^ in eQ, 0Q and 100Q cells. Radiolabeled reaction products were separated by denaturing PAGE alongside a sequencing ladder and were detected using a phosphorimager. Representative data from n=3 independent experiments are shown.

The modified nucleotides in tRNAs play important roles in their function as adaptor molecules during translation^17,18^. Within the anticodon, position 34 (the wobble position) is a hotspot for modified RNA nucleotides that expand the decoding capacity of tRNAs and ensure the fidelity of mRNA codon–anticodon base-pairing. The anticodon-adjacent positions 37 and 38 are often also modified with functional interplays between nucleotides at positions 34 and 37/38 contributing to ensuring the optimal conformation of the anticodon loop (ACL) for codon base-pairing interactions (see for example^19–22^). In some cases, it has been shown that the functional interconnections between different modified nucleotides in the anticodon stem-loop (ASL) extend to include interdependences at the level of tRNA biogenesis, with the presence of particular modified nucleotides being a pre-requisite for the installation of others^23–26^.

Alongside their central role in protein synthesis, tRNAs have also emerged as precursors of tRNA-derived fragments (tsRNAs)^27–30^. While some tRNA-derived fragments may represent long-lived processing intermediates, in other cases, functions of tsRNAs in the regulation of gene expression have been reported. For example, tsRNAs that associate with argonaute proteins and function analogously to microRNAs have been described and others have been implicated in influencing translation by resolving secondary structural elements in mRNAs or associating with translation factors^31–34^. tsRNA levels are often increased upon cellular stress and several stress-sensitive ribonucleases are implicated in the cleavage of tRNAs into tsRNAs, including the RNase A family endoribonuclease angiogenin (ANG), RNase L, RNase T2, Dicer and IRE1α^35–42^. Modified nucleotides present within tsRNAs can influence their biogenesis, stability and functions. For example, the presence of pseudouridine within tRNA fragments containing a 5′ terminal oligoguanine motif influences their association with the polyadenylate binding protein, thus regulating translation initiation^43^. *N*^2^-methylguanosine (m^2^G) or *N*^2,2^-dimethylguanosine (m^2^_2_G) within tsRNAs can enhance their stability, enabling them to exist more persistently^44^. While 5-methoxycarbonyl-methyluridine at position 34 (mcm^5^U_34_) has been suggested to promote tRNA cleavage under certain conditions, 5-methylcytosine (m^5^C), *N*^1^-methyladenosine (m^1^A), ribose 2′-*O*-methylation (Nm) and Q are reported to restrict the cleavage of tRNAs into fragments^45–48^. Although a preventative effect of Q_34_ on tRNA fragmentation has been observed, it has remained unknown whether Q glycosylation contributes to this.

Here, we explore interplays of Q_34_ and its glycosylated derivates with other ACL modifications, define the temporal order in which these modifications are installed and highlight roles of Q glycosylation in modulating sites of ribonucleolytic cleavage and protecting tRNA^Asp(GUC)^ from stress-induced fragmentation.

## RESULTS AND DISCUSSION

### Influence of queuine levels on Q_34_, manQ_34_ and galQ_34_ stoichiometry and other ACL modifications in human tRNA^Asp(GUC)^ and tRNA^Tyr(GUA)^

In tRNA^Asp(GUC)^ and tRNA^Tyr(GUA)^, Q_34_ is inserted by TGT and glycosylated to manQ_34_/galQ_34_ by QTMAN and QTGAL, respectively (**Fig. 1A**). To evaluate the level of queuosinylation and its glycosylated derivatives in tRNA^Asp(GUC)^ and tRNA^Tyr(GUA)^ in HEK293 cells, the modification status of these tRNAs was monitored by acidic gel electrophoresis^49^ and in gels containing *N*-acryloyl-3-aminophenylboronic acid (APB)^50^. In acidic gels, Q/manQ/galQ_34_-containing tRNAs migrate more slowly than tRNAs with G_34_. By contrast, through binding to free *cis*-diol moieties, such as the one present in Q (**Fig. 1A**), the addition of APB during gel electrophoresis specifically retards Q_34_-containing tRNAs relative to G/manQ/galQ_34_-containing tRNAs in which the only free *cis*-diols are those at the 3′-ends of the RNA backbones. RNAs were obtained from cells cultured in the presence of i) fetal bovine serum (FBS), which provides a source of queuine (q) allowing queuosinylation to an endogenous level (eQ), ii) horse serum (HS) that contains only trace amounts of q (0Q) or iii) horse serum supplemented with exogenous q to drive tRNA queuosinylation to its maximal level (100Q). Northern blotting revealed that both tRNA^Asp(GUC)^ and tRNA^Tyr(GUA)^ are sub-stoichiometrically queuosinylated when cultured in FBS (**Fig. 1B, C**). tRNA^Asp(GUC)^ was predominantly queuosinylated whereas the level of G_34_ was markedly higher than Q_34_ in tRNA^Tyr(GUA)^, indicating that tRNA^Asp(GUC)^ is a better substrate of TGT than tRNA^Tyr(GUA)^. Depletion of external q led to the complete loss of (man)Q_34_ in tRNA^Asp(GUC)^ and reduced (gal)Q_34_ in tRNA^Tyr(GUA)^, whereas supplementation with exogenous q fully restored the Q modification in both tRNAs (**Fig. 1B, C**). Comparing the migration patterns of the tRNAs on APB and acidic gels indicated that in eQ cells, the Q-modified tRNAs were predominantly glycosylated, whereas larger fractions of non-glycosylated Q_34_ were detected in tRNA^Asp(GUC)^ and tRNA^Tyr(GUA)^ in 100Q cells, indicating that the levels or rates of mannosylation/ galactosylation are limiting when an excess of q is present.

In *S. pombe*, Q_34_ in tRNA^Asp^ promotes installation of m^5^C at position 38^51^. Bisulfite sequencing was therefore used to monitor the tRNA^Asp(GUC)^-m^5^C_38_ status in eQ, 0Q, and 100Q HEK293 cells. The higher level of (man)Q_34_ in 100Q cells (**Fig. 1B, C**) correlated with increased C_38_ methylation levels (**Fig. 1D**), supporting that stimulation of m^5^C_38_ installation by Q_34_ is conserved from yeast to humans^52^. Notably, in 0Q cells, approximately 70% of tRNA^Asp(GUC)^ contained m^5^C_38_, which may reflect residual Q_34_ present despite growth in media containing only minimal q, but it could also indicate that m^5^C_38_ can be installed independently of Q_34_, albeit less efficiently^51,53^. In tRNA^Tyr(GUA)^, Q_34_ is instead accompanied in the ACL by 1-methylguanosine at position 37 (m^1^G_37_). It was investigated, therefore, whether the extent of queuosinylation affects this methylation. To this end, primer extension assays were performed to detect m^1^G_37_ in tRNA^Tyr(GUA)^ from eQ, 0Q and 100Q cells (**Fig. 1E**). In this assay, nucleotide modifications that lie on the Watson-Crick face, such as methylation of nitrogen 1 of G, perturb reverse transcriptase progression, leading to formation of truncated cDNAs that can be detected by gel electrophoresis. m^1^G_37_ levels were consistent irrespective of the queuosinylation status of the tRNA, implying that this modification is installed independently of Q_34_ (**Fig. 1F**). Interestingly, elevated levels of m^2^_2_ G_26/27_ were observed in 0Q and 100Q cells as compared to eQ cells, supporting an influence of q levels on m^2^_2_ G_26/27_ in HEK293 cells as previously reported **(Fig. 1F)**^54^.

### Bidirectional interdependence of m^5^C_38_ and Q_34_ in tRNA^Asp(GUC)^ and independent installation of m^1^G_37_ and Q_34_ in tRNA^Tyr(GUA)^

As Q_34_ promotes m^5^C_38_ in tRNA^Asp(GUC)^ (**Fig. 1D**), it was investigated whether the cross-talk between these modifications is reciprocal. To this end, a cell line lacking the m^5^C_38_ writer enzyme, DNMT2^55,56^, was established (DNMT2^KO^). A CRISPR guide RNA targeting exon 3 was used to induce a single nucleotide deletion that causes a frameshift leading to the emergence of a premature stop codon 14 codons downstream (p.Gly59Alafs*14; **Fig. 2A**). Lack of DNMT2 in the KO cell line was confirmed by western blotting (**Fig. 2B**) and bisulfite sequencing demonstrated the loss of m^5^C_38_ in tRNA^Asp(GUC)^ (**Fig. 2C**). Analysis of the modification status of position 34 in wild-type and DNMT2^KO^ cells using acidic and ABP gels demonstrated that compared to the sub-stoichiometric queuosinylation observed in wild-type cells, loss of DNMT2 led to complete exchange of G_34_ for (man)Q_34_ (**Fig. 2D, E**). As the APB gel electrophoresis indicated low levels of Q_34_ in both samples, it appears that once installed, Q_34_ is rapidly converted to manQ_34_ and the lack of m^5^C_38_ does not impede the mannosylation step. However, these results also indicate that the presence of m^5^C_38_ limits incorporation of Q_34_ by TGT in cells.

**Figure 2.**
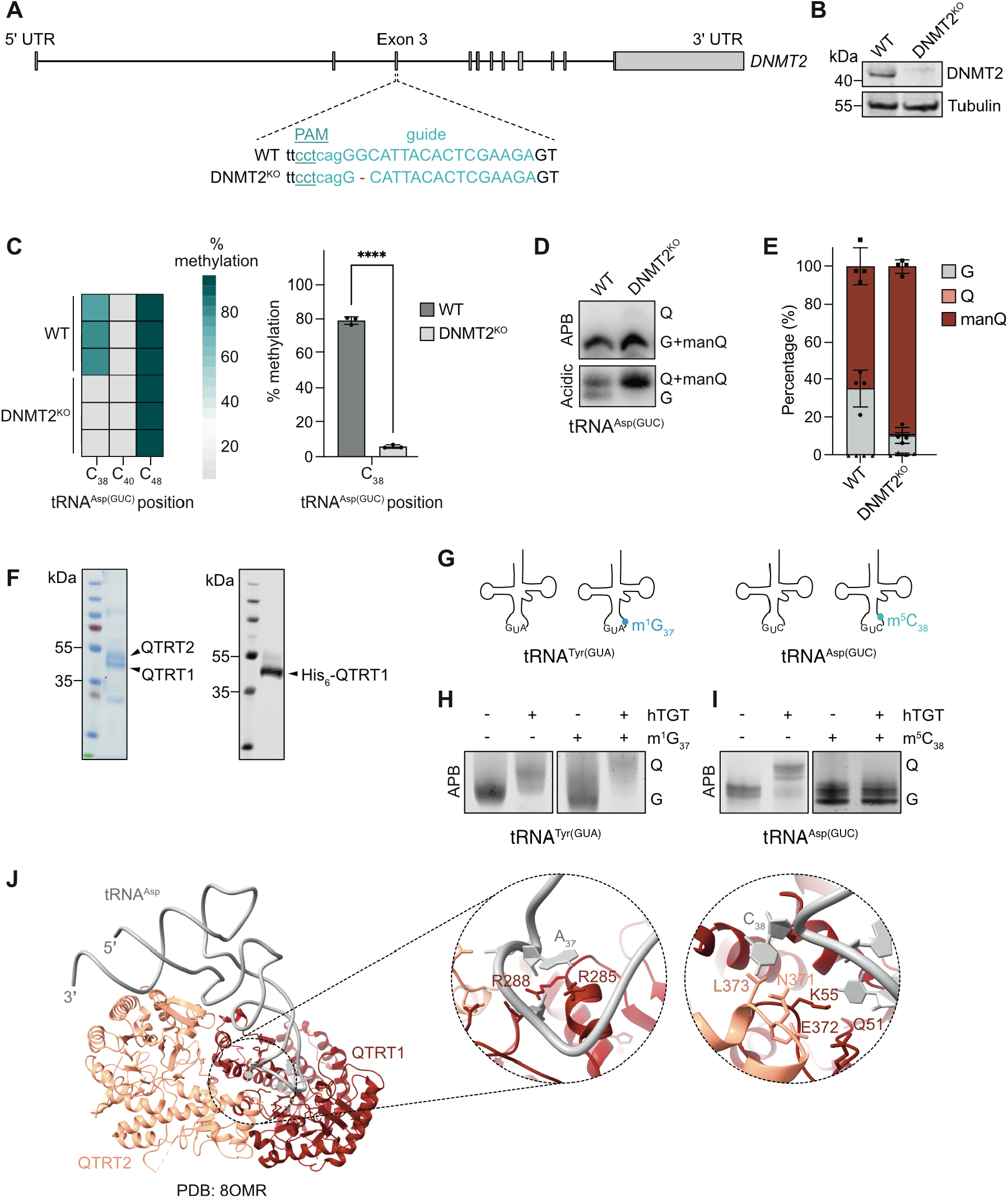
Crosstalk between Q_34_ and tRNA^Asp(GUC)^-m^5^C_38_ and tRNA^Tyr(GUA)^-m^1^G_37_. **(A)** Scheme of *DNMT2* with the genomic region targeted by CRIPSR-Cas9 genome editing highlighted in wild-type (WT) and knockout (DNMT2^KO^) cells. **(B)** Proteins from WT and DNMT2^KO^ cells were analyzed by western blotting using antibodies against DNMT2 and, as a loading control, Tubulin.**(C)** RNAs from WT and DNMT2^KO^ cells were subjected to bisulfite sequencing to monitor m^5^C_38_ levels in tRNA^Asp(GUC)^. Results from n=3 independent experiments are shown as a heatmap (left) and a bar plot (right; error bars represent mean ± standard deviation). **(D)** Total RNAs from WT and DNMT2^KO^ cells were separated by APB (5%) or acidic gel electrophoresis and tRNA^Asp(GUC)^ was detected by northern blotting. Representative image from n=4 independent experiments is shown. **(E)** Quantification of the levels of G_34_, Q_34_ and manQ_34_ in samples as described in (D). Error bars represent mean ± standard deviation. **(F)** QTRT1-His_6_ and QTRT2 were recombinantly co-expressed in *E. coli* and purified. Purified proteins were separated by SDS-PAGE and detected by Coomassie staining (left) and western blotting (anti-His_6_; right). **(G)** Scheme of the modified and unmodified synthetic tRNAs used for queuosinylation assays. **(H, I)** The synthetic tRNAs depicted in (G) were used for *in vitro* queuosinylation assays with purified TGT (F). Reaction products (tRNA^Tyr(GUA)^ (H) and tRNA^Asp(GUC)^ (I)) were separated by ABP (0.5%) gel electrophoresis and RNAs were detected by SYBR Gold staining. **(J)** Model of the human TGT in complex with tRNA^Asp^ (PDB: 8OMR). Magnified views show key residues in the anticodon loop region of the tRNA and the TGT subunits.

To consolidate this notion and explore whether the presence of m^1^G_37_ in tRNA^Tyr(GUA)^ impacts queuosinylation, QTRT1 and QTRT2 were co-expressed in *E. coli* and the TGT complex was purified (**Fig. 2F**). In parallel, synthetic tRNA^Asp(GUC)^ containing C_38_/m^5^C_38_ and tRNA^Tyr(GUA)^ containing G_37_/m^1^G_37_ were prepared (**Fig. 2G; Supplementary Figures S1 and S2**) and *in vitro* queuosinylation assays were performed. Both unmodified and m^1^G_37_-containing tRNA^Tyr(GUA)^ were efficiently queuosinylated (**Fig. 2H**), indicating that the lack of influence of Q_34_ on methylation of G_37_ (**Fig. 1F**) is mirrored by m^1^G_37_-independent installation of Q_34_. The C_38_-containing tRNA^Asp(GUC)^ was, likewise, efficiently queuosinylated by TGT, however, Q_34_ was not installed into tRNA^Asp(GUC)^ containing m^5^C_38_ (**Fig. 2I**). This corroborates the results from DNMT2^KO^ cells, in which queuosinylation and subsequent mannosylation of tRNA^Asp(GUC)^ is favored in the absence of m^5^C_38_, by showing directly that methylation of C_38_ prevents modification of position 34 by TGT.

Analysis of structures of human TGT bound to tRNA^Asp^ or stem loop tRNA substrates^11,12^ offers a mechanistic basis for the inhibitory effect of m^5^C_38_ on Q_34_ installation (**Fig. 2J**). Nucleotides 32 to 35 of the ACL are important for substrate recognition by TGT, while neighboring residues stabilize the tRNA helical stem, potentially optimizing the ACL conformation for catalysis^12,57^. Nucleotides 35 to 38 adopt a zig-zag conformation with residues 36 and 38 flipped out of the loop, and although residue 38 is unlikely to contribute to sequence-specific recognition, it is the only ACL nucleotide contacted by two QTRT2 residues. In the cryo-EM structure^11^, carbon 5 of C_38_ of unmodified tRNA^Asp(GUC)^ is proximal to asparagine 371 of QTRT2 (**Fig. 2J**), suggesting that addition of a methyl group at this position could disrupt contacts with QTRT2, and/or alter ACL conformation. Such effects could impair substrate binding or the structural rearrangements required for Q incorporation, thus rationalizing the observed lack of queuosinylation of m^5^C_38_-containing tRNA^Asp(GUC)^ (**Fig. 2I**). By contrast, residue 37 of the tRNA is oriented towards the interior of the loop and is stabilized through stacking interactions between the base and Arg285 in QTRT1 as well as hydrogen bonding between the O2′ of the ribose and Arg288^12^. Consequently, nitrogen 1 methylation of G_37_ should not interfere with TGT recognition, consistent with efficient Q incorporation into m^1^G_37_-containing substrates (**Fig. 2H**).

### Glycosylations of Q_34_ occur after m^1^G_37_ in tRNA^Tyr(GUA)^ and m^5^C_38_ in tRNA^Asp(GUC)^

The identification of bidirectional crosstalk between m^5^C_38_ and Q_34_ in tRNA^Asp(GUC)^ and the independence of m^1^G_37_ and Q_34_ in tRNA^Tyr(GUA)^ raised the question whether interdependencies exist also at the level of Q_34_ hypermodification to man/galQ_34_. Towards this goal, QTMAN^KO^ and QTGAL^KO^ cell lines were generated. In the case of *QTMAN*, a single nucleotide deletion within exon 5 induced a frame shift (p.Glu103Argfs; **Fig. 3A**), whereas a nucleotide substitution and an insertion within the guide RNA target site in exon 3 of *QTGAL* caused a frameshift that led to a premature stop codon eight amino acids downstream (p.Gly68Trpfs*8; **Fig. 3B**). As QTMAN and QTGAL are expressed at very low levels making antibody-mediated detection of these proteins unattainable, the glycosylation status of Q_34_ of tRNA^Asp(GUC)^ and tRNA^Tyr(GUA)^ was monitored by comparison between the migration patterns of these tRNAs in acidic and ABP gels. The mobility of tRNA^Asp(GUC)^ and tRNA^Tyr(GUA)^ from wild-type and the QTMAN/QTGAL^KO^ in acidic gels indicates that the overall modification status of these tRNAs in not affected (**Fig. 3C**). However, the slower migration in APB gels of tRNA^Asp(GUC)^ from the QTMAN^KO^ cell line and tRNA^Tyr(GUA)^ from the QTGAL^KO^ cell line compared to wild-type indicated the presence of the non-glycosylated forms of Q_34_ in these tRNAs in the KO cells (**Fig. 3C**). The absence of the mannose and galactose moieties in the KO cell lines was further confirmed by treatment with sodium periodate, which oxidizes the *cis*-diol group of Q and therefore eliminates the differential migration of non-glycosylated Q in the APB gels, producing a faster-migrating species (**Fig. 3C**)^49,58^.

**Figure 3.**
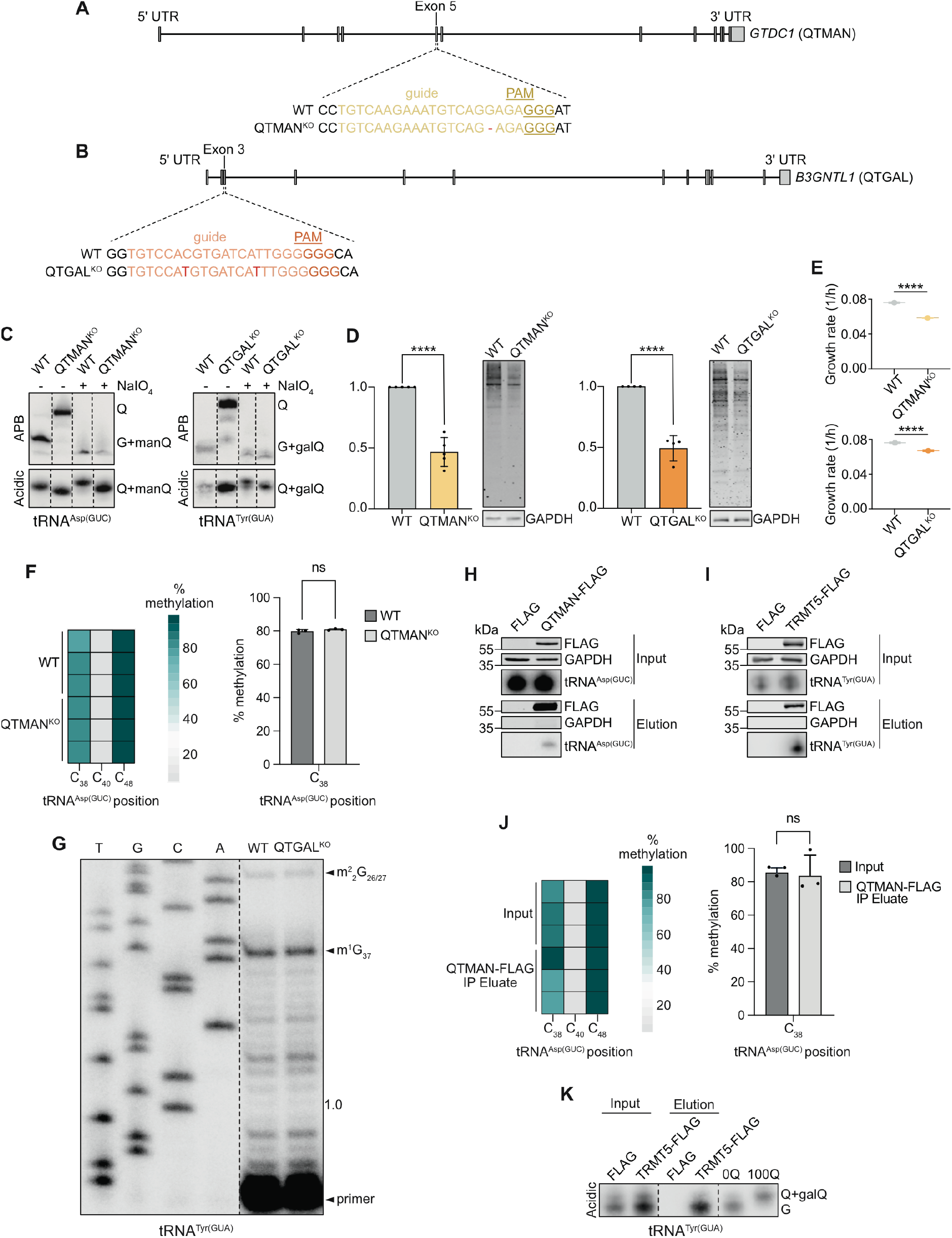
Influence of Q_34_ glycosylations on other ACL modifications. **(A, B)** Schemes of *QTMAN* (A) and *QTGAL* (B) with genomic regions targeted by CRIPSR-Cas9 genome editing highlighted in wild-type (WT) and QTMAN^KO^/QTGAL^KO^ cells. **(C)** Total RNAs from WT and QTMAN^KO^ (left) or QTGAL^KO^ (right) cells were treated or not with sodium periodate (NaIO_4_) and subsequently separated by APB (5%) or acidic gel electrophoresis. tRNA^Asp(GUC)^ (left) or tRNA^Tyr(GUA)^ (right) were detected by northern blotting. **(D)** Nascent proteins in WT and QTMAN/QTGAL^KO^ cells were labeled with AHA. AHA-containing proteins were conjugated to biotin, and total protein was separated by SDS-PAGE and transferred to a membrane. Biotinylated proteins were detected using fluorescently labeled streptavidin, and western blotting for GAPDH served as a loading control. The relative levels of biotinylated proteins were quantified in n=3 independent experiments and are shown as a bar plot (right; error bars represent mean ± standard deviation). **(E)** The growth rates of the wild-type, QTMAN^KO^ (left) and QTGAL^KO^ (right) cell lines were determined by flow cytometry in n=3 independent experiments. **(F)** RNAs from WT and QTMAN^KO^ cells were subjected to bisulfite sequencing to monitor m^5^C_38_ levels in tRNA^Asp(GUC)^. Results from n=3 independent experiments are shown as a heatmap (left) and a bar plot (right; error bars represent mean ± standard deviation). **(G)** Primer extension assay monitoring m^1^G_37_ levels in tRNA^Tyr(GUA)^ in WT and QTGAL^KO^. Radiolabeled reaction products were separated by denaturing PAGE alongside a sequencing ladder and were detected using a phosphorimager. Representative data from n=3 independent experiments are shown. **(H, I)** Extracts from cells expressing QTMAN-FLAG (H), TRMT5-FLAG (I) or the FLAG tag were used for anti-FLAG immunoprecipitation experiments, and proteins and RNAs present in input (0.5%) and eluates were analyzed by western and northern blotting, respectively. **(J)** RNAs from input and eluate samples of immunoprecipitation experiments as in (H) were subjected to bisulfite sequencing to monitor m^5^C_38_ levels in tRNA^Asp(GUC)^. Results from n=3 independent experiments are shown as a heatmap (left) and a bar plot (right; error bars represent mean ± standard deviation). **(K)** RNAs from experiments performed as in (I) were separated by acidic gel electrophoresis and tRNA^Tyr(GUA)^ was detected by northern blotting. Total RNA from 0Q and 100Q cells were used as controls. In (H), (I) and (K), representative data from n=3 independent experiments are shown.

Glycosylation of Q_34_ has previously been demonstrated to optimize translation ^13^ so the QTMAN^KO^/QTGAL^KO^ cell lines were further characterized by analyzing nascent protein synthesis and cellular growth. For quantitative analysis of newly synthesized proteins, wild-type and the QTMAN^KO^/QTGAL^KO^ cells depleted of methionine were grown in the presence of azidohomoalanine (AHA), an analogue of methionine that is accepted by tRNA^Met^ and incorporated into nascent proteins during translation ^59^. Coupling of the incorporated AHA to biotin using click-chemistry enabled their detection with fluorescently-conjugated streptavidin. Decreases in the levels of newly synthesized proteins in the QTMAN/QTGAL^KO^ cell lines as compared to wild-type were observed (**Fig. 3D**), supporting the requirement of Q_34_ glycosylation in tRNA^Asp(GUC)^ and tRNA^Tyr(GUA)^ for efficient translation. Consistent with the importance of protein synthesis for cell proliferation, growth of the KO cell lines was perturbed relative to wild-type cells (**Fig. 3E**).

The influence of Q_34_ glycosylation on the other ACL modifications in tRNA^Asp(GUC)^ and tRNA^Tyr(GUA)^ was then examined. To this end, bisulfite sequencing was used to monitor m^5^C_38_ levels in tRNA^Asp(GUC)^ from wild-type and QTMAN^KO^ cells. No difference in the methylation levels of C_38_ was observed upon loss of Q_34_ mannosylation, indicating that the presence of the mannose moiety does not interfere with DNMT2 activity (**Fig. 3F**). Similarly, primer extension assays (**Fig. 1E**) demonstrated that lack of galactosylation of Q_34_ in tRNA^Tyr(GUA)^ does not impact methylation of G_37_ by TRMT5 (**Fig. 3G**). To gain insight into the order in which tRNA^Asp(GUC)^ and tRNA^Tyr(GUA)^ are methylated and glycosylated, the modification statuses of position 38 of QTMAN-bound tRNA^Asp(GUC)^ and position 34 of tRNA^Tyr(GUA)^ associated with the m^1^G_37_ methyltransferase TRMT5 were assessed. Immunoprecipitation of QTMAN-FLAG or TRMT5-FLAG enriched tRNA^Asp(GUC)^ and tRNA^Tyr(GUA)^, respectively (**Fig. 3H, I**). Monitoring C_38_ methylation in tRNA^Asp(GUC)^ bound to QTMAN revealed a similar level of m^5^C to that present in total tRNAs, indicating that methylation by DNMT2 precedes Q_34_ mannoslyation (**Fig. 3J**). ABP/acidic gel electrophoresis of tRNA^Tyr(GUA)^ recovered with TRMT5 revealed that predominantly G_34_-containing tRNA was present (**Fig. 3K**). This suggests that TRMT5 acts on tRNA^Tyr(GUA)^ prior to TGT and QTGAL, further supporting the structure-based notion that methylation of G_37_ does not affect tRNA binding by TGT (**Fig. 2J**), and suggesting that this is also the case for QTGAL.

### Q_34_ glycosylation limits tRNA fragmentation in response to oxidative stress

Non-glycosylated Q_34_ is implicated in limiting fragmentation of tRNA^Asn/His^ in cells^46^, so the impact of Q_34_, manQ_34_ and galQ_34_ on stress-induced cleavage of tRNA^Asp(GUC)^ and tRNA ^Tyr(GUA)^ was explored. The RNase A family endoribonuclease, ANG, plays a central role in the cleavage of tRNAs within the ACL during stress so cleavage assays were performed with recombinant ANG and *in vitro* transcribed tRNA^Asp(GUC)^. Compared to other RNase A family endoribonucleases, the C-terminal region of ANG occludes the catalytic site^60^ but association with stalled ribosomes relieves this auto-repression^38^. Substitution of glutamine 117 for glycine (Q117G)^60^ similarly repositions the C-terminal region of ANG to unblock the active site, rendering it an effective mimic of the activated form of the enzyme (**Fig. 4A**). For *in vitro* tRNA cleavage assays, wild-type ANG, ANG_Q117G_ and ANG_Q117G_ in which histidine 114 within the catalytic core was substituted for alanine (ANG_H114A+Q117G_) to block catalytic activity were overexpressed in *E. coli* and purified (**Fig. 4B**). Cleavage assays demonstrated increased activity of ANG_Q117G_ compared to wild-type as well as the effectiveness of the H114A substitution in blocking ANG catalytic activity (**Fig. 4C**).

**Figure 4.**
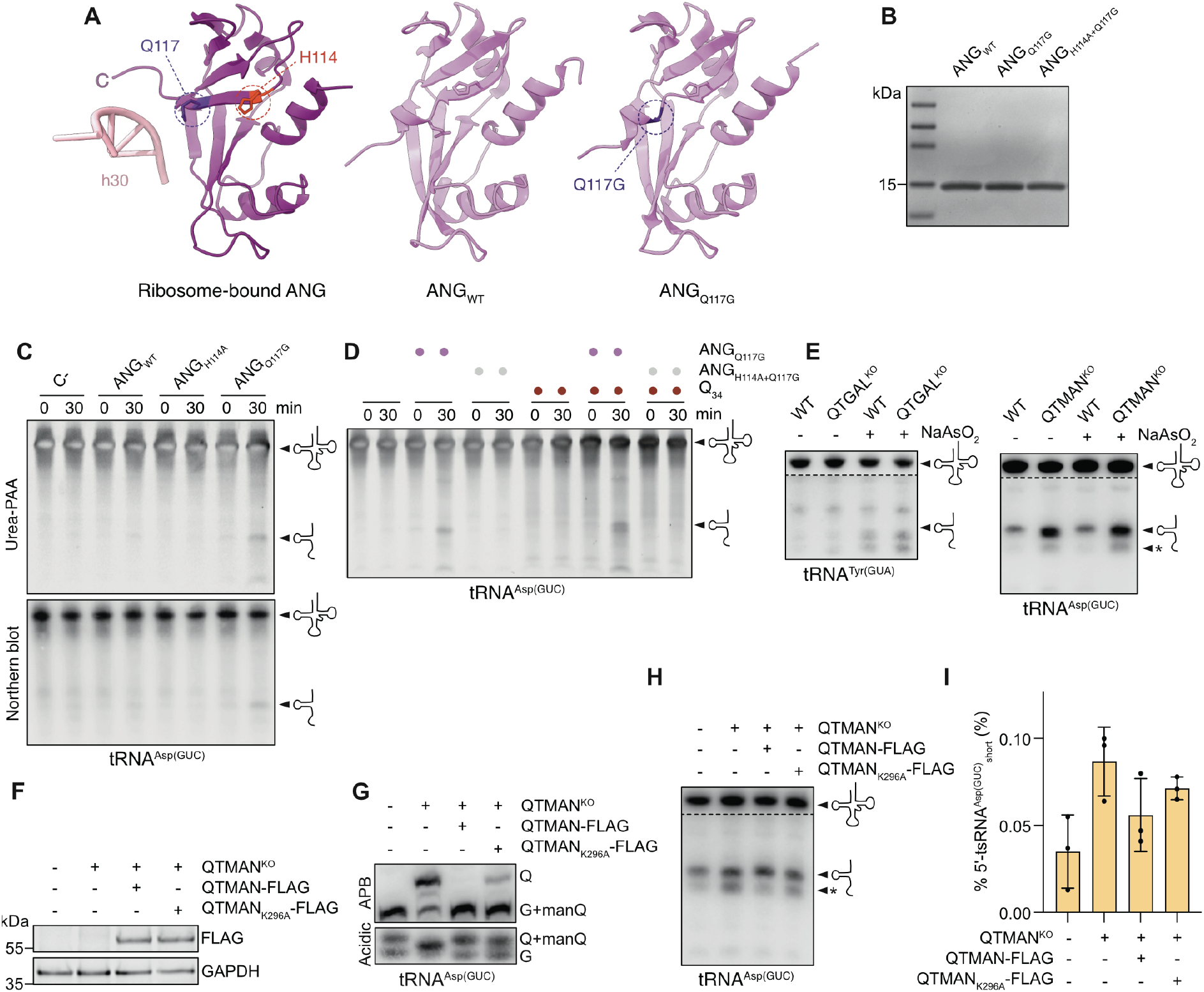
tRNA fragmentation in QTMAN^KO^ and QTGAL^KO^ cells. **(A)** Structures of ribosome-bound ANG (left) (PDB: 9BDL), ANG_WT_ (middle) (PDB: 1ANG) and ANG_Q117G_ (right) (PDB: 1K59). H114 and Q117G residues are highlighted on the structures. **(B)** ANG_WT_, ANG_Q117G_ and ANG_H114A/Q117G_ were recombinantly expressed in *E. coli* and purified. Purified proteins were separated by SDS-PAGE and detected by Coomassie staining. **(C)** *In vitro* transcribed tRNA^Asp(GUC)^ and the recombinant ANG proteins (A) were used for *in vitro* cleavage assays. C-indicates samples to which no nuclease was added. Reaction products were separated by denaturing gel electrophoresis and detected by SYBR Gold staining and northern blotting. **(D)** *In vitro* tRNA cleavage assays were performed as in (C) using *in vitro* transcribed tRNA^Asp(GUC)^ containing or not Q_34_. Reaction products were separated by denaturing gel electrophoresis and detected by SYBR Gold staining. Representative data from n=3 independent experiments are shown. **(E)** Wild-type (WT) and QTMAN^KO^ (left) and QTGAL^KO^ (right) cells were treated or not with sodium arsenite (NaAsO_2_) to induce oxidative stress. Total RNAs were extracted and northern blotting for tRNA^Asp(GUC)^ (left) and tRNA^Tyr(GUA)^ (right) enabled the detection of full-length tRNAs (upper) and tRNA-derived fragments (lower). The asterisk (*) indicates the additional shorter tRNA fragment species (5’-tsRNA^Asp(GUC)^_short_). Representative data from n=3 independent experiments are shown. Note that regions above and below the dotted line represent different exposures of the same membrane. **(F)** Proteins from WT and QTMAN^KO^ cells, and QTMAN^KO^ cells expressing QTMAN-FLAG or QTMAN_K296A_-FLAG, were analyzed by western blotting using antibodies against the FLAG-tag and, as a loading control, GAPDH. **(G)** RNAs from cells as described in (F) were separated by APB (5%) or acidic gel electrophoresis and tRNA^Asp(GUC)^ was detected by northern blotting. Representative data from n=3 independent experiments are shown. **(H)** Cells as in (F) were subjected to oxidative stress, and tRNA^Asp(GUC)^ and tRNA^Asp(GUC)^-derived fragments were detected by northern blotting. The asterisk (*) indicates the additional shorter tRNA fragment species (5’-tsRNA^Asp(GUC)^_short_). Representative data from n=3 independent experiments are shown. **(I)** Quantification of the levels of 5′-tsRNA^Asp(GUC)^_short_ in samples as described in (H). Error bars represent mean ± standard deviation.

The influence of Q_34_ on tRNA^Asp(GUC)^ cleavage by ANG was then tested. Q_34_ was stoichiometrically installed in *in vitro* transcribed tRNA^Asp(GUC)^ using recombinant TGT (**Fig. 2F**) and ANG-mediated cleavage assays were performed using equal amounts of modified and unmodified substrates. Strikingly, cleavage products of both the G_34_- and Q_34_-containing transcripts were detected at similar levels (**Fig. 4D**), suggesting that Q_34_ alone does not prevent fragmentation of these tRNAs. To evaluate whether glycosylation of Q_34_ influences tRNA cleavage, fragmentation of tRNA^Asp(GUC)^ and tRNA^Tyr(GUA)^ from wild-type and QTMAN/QTGAL^KO^ cells was examined in non-stressed cells and those exposed to sodium arsenite-induced oxidative stress. Northern blotting revealed elevated levels of tRNA^Tyr(GUA)^-derived fragments in stressed compared to unstressed cells (**Fig. 4E; left**). In stressed cells, loss of galactosylation of Q_34_ led to a very modest accumulation of specific fragments of tRNA^Tyr(GUA)^ (**Fig. 4E; left**), implying that Q_34_ galactosylation only minimally contributes to limiting tRNA cleavage. In the absence of oxidative stress, a single fragment of tRNA^Asp(GUC)^ was detected in wild-type cells, whereas in the QTMAN^KO^ cell line, this fragment not only accumulated more strongly, but also an additional shorter fragment of tRNA^Asp(GUC)^ (5′-tsRNA^Asp(GUC)^_short_) was detectable (**Fig. 4E; right**), indicating that mannosylation of Q_34_ inhibits tRNA cleavage at specific sites. Together, these data demonstrate that Q_34_ mannosylation can influence the site of tRNA^Asp(GUC)^ cleavage and contribute to the protection of this tRNA against fragmentation.

As loss of Q_34_ mannosylation in tRNA^Asp(GUC)^ had a pronounced effect on tRNA fragmentation, a complementation system was established in which FLAG-tagged, wild-type QTMAN or a version in which lysine 296 that is proposed to contribute to the catalytic reaction^13^ was substituted for alanine (K296A) (**Fig. 4F**). To demonstrate the requirement of lysine 296 for formation of manQ_34_, the modification status of tRNA^Asp(GUC)^ from these cell lines was assessed. Consistent with the previous result (**Fig. 3C**), Q_34_, rather than manQ_34_, was detected in the QTMAN^KO^ cell line (**Fig. 4G**). Re-expression of QTMAN-FLAG, but not QTMAN_K296A_-FLAG, restored the manQ_34_ level to that of the wild-type cells (**Fig. 4G**), confirming the importance of lysine 296 for the mannosylation reaction. Upon oxidative stress, northern blotting for tRNA^Asp(GUC)^ demonstrated that expression of QTMAN-FLAG in the QTMAN^KO^ background reduced the levels of the 5′-tsRNA^Asp(GUC)^_short_ fragment similar to the endogenous level (**Fig. 4H, I**). A similar pattern of tRNA^Asp(GUC)^-derived fragments to that observed in the QTMAN^KO^ cell line was detected in cells expressing QTMAN_K296A_-FLAG (**Fig. 4H, I**), confirming that the increased tRNA cleavage at additional site arises due to loss of the Q_34_ mannosylation.

Taken together, the data presented here demonstrate a modification pathway of the human tRNA^Asp(GUC)^ ACL in which queuosinylation of G_34_ by TGT to produce Q_34_ favors installation of m^5^C_38_ by DNMT2. The presence of m^5^C_38_ inhibits the action of TGT such that any tRNAs methylated prior to association with TGT are not queuosinylated (**Fig. 5A**). The sequential incorporation of these RNA modifications, together with their bidirectional effects on the corresponding modification enzymes, can ensure appropriate modification levels and constitute a feedback mechanism to dynamically fine-tune the ACL modification status of tRNA^Asp(GUC)^ to optimize its function. Q_34_ hypermodification occurs after m^5^C_38_ installation (**Fig. 5A**) but methylation of C_38_ does not impact mannosylation, implying that substrate recognition by QTMAN does not involve sensing the prior modification. Although Q_34_ installation in tRNA^Tyr(GUA)^ can occur independently of m^1^G_37_, our results suggest hierarchical installation of these modifications in cells, with G_37_ being methylated by TRMT5 prior to installation of Q_34_ by TGT (**Fig. 5B**). As queuosinylation has recently been shown to precede tRNA^Tyr(GUA)^ splicing^16^, this implies that introduction of m^1^G_37_ by nuclear TRMT5^61^ is among the earliest tRNA maturation events. tRNA fragments are consolidating as an important class of noncoding RNAs, emphasizing the relevance of understanding their origins and identifying factors that influence their production. Alongside its function in facilitating efficient decoding^13^, our results highlight an additional role of manQ_34_ in tRNA^Asp(GUC)^ in determining sites of tRNA cleavage and suppressing tRNA fragmentation (**Fig. 5A**).

**Figure 5.**
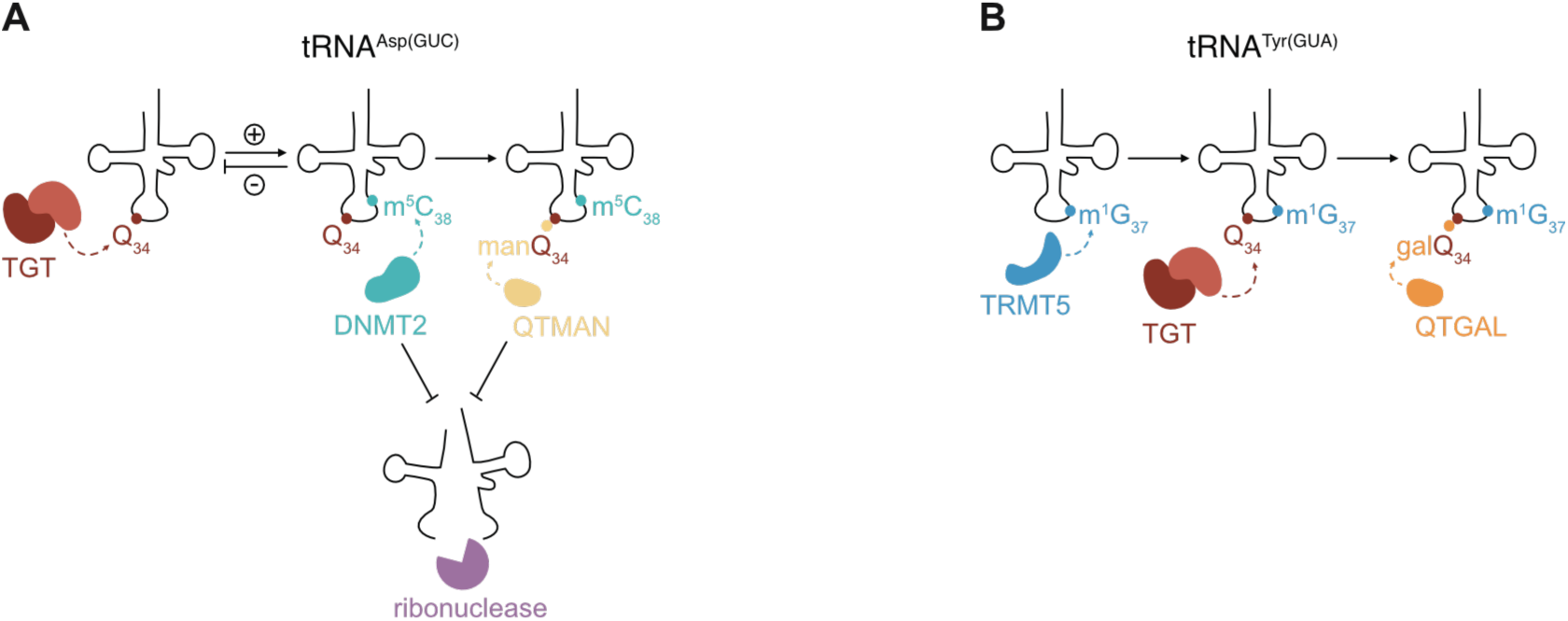
Model of interplays between man/galQ_34_ in tRNA^Asp(GUC)/Tyr(GUA)^ and other ACL modifications and the role of Q_34_ glycosylation in limiting tRNA fragmentation. **(A)** In tRNA^Asp(GUC)^, queuosinylation by TGT to produce Q_34_ stimulates the methyltransferase DNMT2 to methylate C_38_ (m^5^C_38_). The presence of m^5^C_38_ inhibits TGT activity, demonstrating a specific order of installation for these modifications. The queuosinylated and methylated tRNA is subsequently mannosylated by QTMAN. The presence of m^5^C_38_^45^ and/or manQ_34_ limits tRNA^Asp(GUC)^ fragmentation. **(B)** In tRNA^Tyr(GUA)^, G_37_ residue is likely methylated by TRMT5 to form m^1^G_37_ prior to queuosinylation by TGT, which is then followed by Q galactosylation mediated by QTGAL to form galQ_34_.

## MATERIALS AND METHODS

### Human cell culture

HEK293 Flp-In T-REx (ThermoFisher Scientific) cells were cultured in Dulbecco’s Modified Eagle Medium (DMEM; ThermoFisher Scientific) supplemented with 1% (v/v) penicillin-streptomycin (ThermoFisher Scientific) and 10% (v/v) fetal bovine serum (FBS; Sigma-Aldrich). Cells were cultured at 37 °C in a humidified atmosphere with 5% CO_2_ and passaged at a ratio of 1:10 by trypsinization using 0.25% Trypsin-EDTA (ThermoFisher Scientific). For generation of 0Q cells^46,49^, HEK293 Flp-In T-REx cells were cultured as usual, but replacing FBS with 10% horse serum (HS; ThermoFisher Scientific), which contains only trace amounts of q^62–65^. Cells were passaged at least 10 times under these conditions to ensure depletion of Q modification^46,49^. To generate 100Q cells, 0Q cells were grown to 70-80% confluency and 1 µM queuine (Toronto Research) was added for 24 h to enable tRNAs to become fully Q-modified. Oxidative stress was induced by addition of 500 µM sodium (meta)arsenite (NaAsO_2_) (Sigma-Aldrich) for 1 h.

### Generation of KO cell lines by CRIPSR-Cas9 genome editing

Sequences encoding single guide RNAs (sgRNAs) (**Supplementary Table S1**) targeting selected genomic regions were assembled by annealing oligonucleotides and phosphorylating them using T4 polynucleotide kinase (PNK; ThermoFisher Scientific). Each oligonucleotide duplex was subsequently cloned into the px459 plasmid that enables expression of sgRNAs from the U6 promoter as well as the Cas9 endonuclease^66^. HEK293 Flp-In T-REx cells were transfected with the px459 plasmids using X-tremeGENE 9 DNA Transfection Reagent (Roche) according to the manufacturer’s instructions. Transfected cells were selected with 1 µg/mL puromycin (Sigma-Aldrich) for 7-10 days. Genomic cleavage efficiency was tested with the GeneArt Genomic Cleavage Detection Kit (ThermoFisher Scientific) according to manufacturer’s instructions. Cells were then seeded at single cell density in 96-well plates and expanded. Genomic DNA (gDNA) was extracted using the PureLink Genomic DNA kit (ThermoFisher Scientific) and the extracted gDNA was used as a template to amplify the targeted genomic region via polymerase chain reaction (PCR) using the oligonucleotide primers listed in **Supplementary Table S2**. Amplicons were analyzed by Sanger sequencing to confirm presence of genomic variations.

### Generation of stably transfected cell lines using the Flp-In system

The coding sequences of *TRMT5* (NM_020810.3) or *QTMAN* (NM_001006636.5) were cloned into a pcDNA5-based plasmid for the expression of C-terminally His_6_-2×FLAG (FLAG)-tagged proteins (**Supplementary Table S3**). Site-directed mutagenesis was used to introduce mutations leading to the substitution of lysine 296 within QTMAN for alanine (K296A) with the oligonucleotides listed in **Supplementary Table S4**. HEK293 Flp-In T-REx cells or the QTMAN^KO^ cell line were co-transfected with the pOG44 plasmid encoding the Flp recombinase and the appropriate pcDNA5-based plasmid. Transfections were performed using the X-tremeGENE 9 DNA Transfection Reagent (Roche) following the manufacturer’s instructions. After 48 h, selection of cells in which the *trans*gene had been integrated into the Flp-In locus were selected with 100 µg/mL Hygromycin B (AppliChem) and 10 µg/mL Blasticidin S (Carl ROTH). Selection was carried out for 14 days. To induce expression of the *trans*genes, cells were treated with 1 µg/mL (TRMT5-FLAG) or 2 ng/mL (QTMAN-FLAG/QTMAN_K296A_-FLAG) tetracycline (Sigma-Aldrich) for 24 h or 72 h.

### Cell counting by flow cytometry

Cell counting by analytical flow cytometry was performed in a FACSCant II (BD Biosciences) using CountBright Plus Absolute Counting Beads (ThermoFisher Scientific). Wild-type and QTMAN/ QTGAL^KO^ cells were seeded in 24-well plates at a final concentration of 1,000 cells/well. After 12 h, first measurements were obtained by staining attached cells with Hoechst 34580 (ThermoFisher Scientific) at a final concentration of 2 µg/mL for 30 min at 37 °C. Cells were harvested and incubated with 10 or 25 µL of absolute counting beads before counting. Data acquisition and gating were performed by using the FACS Diva software (v.9.2) and a total of 2,500 bead events were collected for each sample. Cell staining and counting procedures were performed every 24 h to 96 h after the first measurement. Data were analyzed in R software (v.4.1.2) using the package flowCore (v.2.6.0). The absolute count (concentration of cells in the original sample) was determined using a custom-made R script based on the ratio of the cell volume collected and the counting beads added to the samples as described by the manufacturer using the following equation:

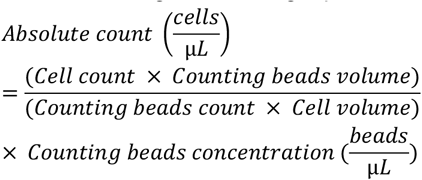

The absolute count of each sample was normalized to the first measurement of each replicate. Normalized data was further analyzed using GraphPad Prism (v.10.2.2) by performing a non-linear regression and data was fitted to an exponential (Malthusian) growth model constraining the initial value Y_0_ to 1.

### Total protein extraction and quantification, and western blotting

Cell pellets were resuspended in RIPA buffer (150 mM NaCl, 1% NP-40, 0.5% sodium deoxycholate, 0.1% (w/v) SDS, 50 mM Tris pH 8.0, 5 mM MgCl_2_), incubated on ice for 30 min, and centrifugated at 20,000 xg for 5 min. The supernatant containing the total protein extract was quantified using the Pierce BCA Protein Assay kit (ThermoFisher Scientific).

SDS-polyacrylamide gel electrophoresis (SDS-PAGE) was performed by using either Tris-HCl (Laemmli) or Bis-Tris gel and buffer systems. For western blotting analyses, samples were transferred onto Amersham Protran nitrocellulose membranes (0.45 µm pore size; Cytiva) using a wet-transfer system. Membranes were blocked for 1 h at room temperature in Tris-buffered saline (TBS) supplemented with 0.1% Tween 20 (TBST) and 5% (w/v) milk powder and incubated with primary antibodies (**Supplementary Table S5**) diluted in TBST+5% milk powder for 16 h at 4 °C. Membranes were then washed thoroughly and incubated with secondary antibodies (**Supplementary Table S6**) diluted in TBST+5% milk powder for 1 h at room temperature. Membranes were further washed and signals were detected using an Odyssey-CLx (LICORbio). Analyses and quantifications were performed in ImageStudio Lite software (LICORbio).

### AHA-labeling to monitor nascent protein synthesis

Quantitative analysis of newly synthesized proteins was performed by using L-azidohomoalanine (AHA)-labeling^59,67,68^. Wild-type and QTMAN/QTGAL^KO^ cells were grown in methionine-free DMEM (ThermoFisher Scientific) supplemented with 10% FBS for 1 h. 100 µM L-azidohomoalanine (Tocris Bioscience) was added and the cells were incubated for a further 3 h. Cells were harvested, resuspended in AHA Lysis Buffer (50 mM Tris-HCl pH 8.0; 0.5% SDS) and lysed by sonication in a Bandelin Electronic Sonopuls HD2070 device. After centrifugation at 20,000 xg for 15 min at 4°C, soluble lysates containing 90 µg of total protein were incubated with 100 µM Biotin-Alkyne (Sigma-Aldrich), 100 mM Tris(2-carboxyethyl)phosphine (TCEP; Sigma-Aldrich), 100 µM Tris(3-hydroxypropyltriazolylmethyl)amine (THPTA; Sigma-Aldrich), and 1 mM CuSO_4_ (Sigma-Aldrich). After 1 h at room temperature, proteins were precipitated with trichloroacetic acid (TCA). Protein pellets were resuspended in NuPAGE LDS Sample Buffer (4x) (ThermoFisher Scientific), separated by Bis-Tris gels, and transferred to nitrocellulose membranes. Membranes were analyzed by western blotting using loading control primary antibodies (**Supplementary Table S5**) as well as fluorescently-labelled streptavidin (IRDye 800CW Streptavidin, LICORbio, 1:10,000). Signals were detected using an Odyssey-CLx (LICORbio). The streptavidin signals of each lane were quantified, normalized according to the GAPDH loading control, and the signals obtained in the QTMAN^KO^ and QTGAL^KO^ lanes were compared to the wild-type samples to obtain the relative fluorescence. Analyses and quantifications were performed in ImageStudio Lite software (LICORbio).

### RNA extraction, denaturing PAGE and northern blotting

Total RNA was extracted from cells or liquid samples using TRI Reagent or TRI Reagent LS (Sigma-Aldrich) following the manufacturer’s instructions. RNAs <200 nucleotides were enriched using the *mir*Vana miRNA Isolation kit (ThermoFisher Scientific) according to the manufacturer’s instructions.

RNA samples in RNA Loading Dye (80% formamide, 1 mM EDTA, 0.025% bromophenol blue; 0.025% xylene cyanol) were denatured at 75 °C for 5 min and separated on 12% denaturing (7 M urea) polyacrylamide Tris-borate-EDTA (TBE) gels. RNAs were transferred to Hybond-N nylon membranes (GE Healthcare) by electroblotting, before crosslinking to the membranes by incubation with 160 mM 1-ethyl-3-(3-dimethylaminopropyl) carbodiimide (EDC) (Sigma-Aldrich) in 130 mM 1-methylimidazole (Sigma-Aldrich), pH 8.0 at 50 °C for 2 h. Membranes were pre-hybridized in SES1 buffer (250 mM sodium phosphate pH 7.0, 7% SDS, 1 mM EDTA) for 30 min at 37 °C. For detection of target RNAs, 5′-radiolabeled probes were generated by incubating 20 pmol DNA oligonucleotide, 20 µCi γ-[^32^P]-ATP (PerkinElmer / Hartmann Analytic) and 10 U T4 PNK at 37 °C for 40 min. The [^32^P]-labeled DNA oligonucleotides (**Supplementary Table S7**) were diluted in SES1 and added to the membranes at 37 °C for 16 h. Membranes were subsequently washed with 6x SSC (50 mM NaCl, 15 mM sodium citrate) followed by 2× SSC + 0.1% SDS each for 30 min at 37 °C. Membranes were exposed to phosphorimager screens and radioactive signals were detected with a Typhoon FLA 9500 scanner (GE Healthcare). Images were analyzed and quantified in ImageStudio Lite software (LICORbio).

### ABP gel electrophoresis

Total RNA samples (2.5 µg) were deacylated by incubation in 100 mM Tris-HCl pH 9.0 with 20 U RiboLock RNase Inhibitor at 37 °C for 30 min. Samples were mixed with an equal volume of APB Loading Dye (2×) (7 M urea, 100 mM EDTA, 0.05% bromophenol blue, 0.05% xylene cyanol) and separated on 10% denaturing (7 M urea) polyacrylamide TAE gels supplemented with 0.5% or 5% *N*-acryloyl-3-aminophenylboronic acid (APB) (Frontier Scientific). RNAs were stained with SYBR Gold Nucleic Acid Gel Stain (ThermoFisher Scientific) and detected using a Sapphire FL Biomolecular Imager (Azure Biosystems) or were transferred to Hybond-N nylon membranes (GE Healthcare) in 0.5x TAE. RNAs were crosslinked to membranes by irradiating twice with UV light at 254 nm at 120 mJ/cm^2^ in a Stratalinker UV Crosslinker (Stratagene) and detected by northern blotting.

### Acidic denaturing gel electrophoresis

Total RNA samples (2.5 µg) were deacylated, mixed with equal volume of acidic RNA Loading Dye (2×) (7 M urea, 170 mM acetate buffer pH 4.8, 0.05% bromophenol blue, 0.05% xylene cyanol) and separated on 12% acid denaturing (7 M urea, 170 mM acetate pH 4.8) polyacrylamide TAE gels. Electrophoresis was performed in TAE running buffer (40 mM Tris, 20 mM acetic acid, 1 mM EDTA) supplemented with 0.17 M acetate buffer pH 4.8 and RNAs were subsequently transferred to Hybond-N nylon membranes (GE Healthcare) in 0.5x TAE. Crosslinking was performed using UV light and tRNAs were detected by northern blotting.

### Quantification of G_34_, Q_34_ and man/galQ_34_ levels using ABP and acidic gel electrophoresis and quantification of 5’-tsRNA levels

The levels of G_34_ and Q_34_ were obtained by quantifying the lower band of the acidic gels and the upper band of APB gels (respectively), whereas the glycoQ_34_ levels were defined as the reminiscent percentage from the total. The percentages of 5′-tsRNAs were obtained by quantifying the corresponding band of the fragment in the northern blots and normalizing by the quantification of the full-length tRNA in the respective lane. Images were analyzed and quantified in ImageStudio Lite software (LICORbio).

### Periodate treatment

Total RNA samples were deacylated and concentrated using the Zymo RNA Clean & Concentrator-5 kit (Zymo Research) following manufacturer’s instructions. Concentrated RNA samples (2.5 µg) were transferred to fresh tubes and incubated with Periodate Mix (final concentration 50 mM sodium (meta)periodate (Sigma-Aldrich); 0.1 M acetate buffer pH 4.8) for 30 min at room temperature. Reactions were quenched with 0.17 M ribose for 5 min at room temperature and mixed with equal volume of APB or Acidic RNA Loading Dye (2×).

### Primer extension

Total RNA (2 µg) was annealed to 0.025 pmol of [^32^P]-labelled oligonucleotide primer (**Supplementary Table S8**) in 1x FS Buffer (ThermoFisher Scientific) by denaturing the mixture at 95°C for 3 min and subsequently cooling down to 55°C. Extension reactions were performed in 1x FS Buffer by adding 100 U SuperScript III Reverse Transcriptase (ThermoFisher Scientific), 1.6 nmol dNTP mix, and 20 U RiboLock RNase Inhibitor (ThermoFisher Scientific) and incubating at 50°C for 30 min. Reactions were quenched with RNA Loading Dye and samples were separated on 15% denaturing (7 M urea) polyacrylamide gels in TBE buffer (89 mM Tris-HCl pH 8.3, 89 mM boric acid, 2 mM EDTA) alongside a sequencing ladder generated by using the Thermo Sequenase Cycle Sequencing kit (ThermoFisher Scientific) according to manufacturer’s instructions using a tRNA^Tyr(GUA)^ PCR product as template. Gels were dried, and exposed to phosphorimager screens (GE Healthcare) before detection of radioactive signals using a Typhoon FLA 9500 scanner (GE Healthcare).

### Bisulfite sequencing for site-specific detection of m^5^C

Total RNA (6 µg) was treated with 4 U TURBO DNase (ThermoFisher Scientific) at 37 °C for 30 min before re-extraction using phenol:chloroform:isoamylalcohol (24:25:1) and ethanol precipitation. RNAs were resuspended in nuclease-free water and bisulfite treatment was performed by using the EpiTect Bisulfite kit (Qiagen) according to manufacturer’s instructions.

Deaminated samples were purified on Micro Bio-Spin P-6 Gel Columns in 6x SSC Buffer (Bio-Rad). For desulfonation, 1 M Tris-HCl pH 9.0 was mixed at an equal volume with samples and incubated at 37 °C for 30 min. Samples were ethanol precipitated and the desulfonated RNA was reverse transcribed with SuperScript III Reverse Transcriptase for cDNA synthesis according to the manufacturer’s instructions using an oligonucleotide primer for tRNA^Asp(GUC)^ (**Supplementary Table S9**). cDNAs were amplified by PCR with KOD One PCR Master Mix (Sigma-Aldrich) using oligonucleotide primers (**Supplementary Table S9**) to add 5′ and 3′ adaptor sequences. PCR products were agarose gel purified using the NucleoSpin Gel and PCR Clean-up kit (Macherey-Nagel) following the manufacturer’s instructions. Eluted samples were used as templates for another PCR reaction with oligonucleotide primers designed for addition of Illumina adapters with specific indexes for multiplexing (TruSeq small RNA). Final PCR products were purified using AMPure XP Beads (Beckman Coulter) and eluates were separated on 3% agarose gels in 1x TBE. Fragments of interest were excised and purified using MinElute Gel Extraction kit (Qiagen) following manufacturer’s instructions. Libraries were quantified by using the Qubit dsDNA High Sensitivity Assay Kit (ThermoFisher Scientific) following manufacturer’s instructions and pooled. Libraries were sequenced on Novaseq6000 at the NGS Integrative Genomics Core Unit (University Medical Centre Göttingen). Quality control analysis of data was performed using FASTQC (v.0.11.5) and paired-end FASTQ files were assembled using PEAR (v0.9.6) with the following parameters: -p 0.05 -j 23 -n 4 -v 6. Due to library multiplexing during sequencing, R2 was used as the forward-oriented read to reconstruct the expected amplicon sequence. Assembled reads were then filtered based on the expected PCR product length using custom Bash scripts implemented with awk (v5.0.1). The 5-nucleotide random barcode (NNNNN) and additional five nucleotides at the 5’ end were trimmed using Cutadapt (v4.9) with the settings -u 10 -j 4. The resulting FASTQ files were again assessed for base composition with FASTQC and quality metrics were summarized using MultiQC (v1.13.dev0). To calculate methylation percentages, trimmed FASTQ reads were analyzed using Bismark v0.24.2. The mature human tRNA-Asp(GUC) sequence was retrieved in FASTA format from the High Confidence Mature tRNA Sequences dataset in GtRNAdb for GRCh38 and used as the reference for Bismark genome preparation. Reads were aligned to the indexed tRNA-Asp(GUC) reference, and methylation levels were determined using the *bismark_methylation_extractor* module with the settings --bedGraph and --cytosine_report. The resulting methylation calls were further processed and visualized in R using custom analysis scripts.

### Immunoprecipitation (IP)

Stable HEK293 Flp-In T-REx cell lines carrying the FLAG or TRMT5-FLAG insertion were cultivated in 15 cm^2^ dishes and induced with 1 µg/mL tetracycline for 24h. Cell pellets were resuspended in TMN150 buffer (50 mM Tris-HCl pH 7.8, 150 mM NaCl, 1.5 mM MgCl_2_, 0.1% NP-40, 5 mM b-mercaptoethanol) supplemented with cOmplete EDTA-free mini protease inhibitor cocktail (Roche) and sonicated in a Bandelin Electronic Sonopuls HD2070 device. Input samples (1%) were transferred to new tubes and the remaining supernatants were incubated with anti-FLAG M2 magnetic beads (Sigma-Aldrich) equilibrated in TMN150 buffer for 2 h at 4°C with mild rotation. Beads were washed with TMN150 buffer and samples were eluted twice with 150 μg/ml 3x FLAG peptide solution (Sigma-Aldrich) at 4°C for 30 min with mild rotation. Eluates were pooled for subsequent protein precipitation and RNA extraction. Proteins were precipitated with TCA and analyzed by western blotting. Total RNA extracted using the TRI Reagent LS (Sigma-Aldrich) and was analyzed by denaturing PAGE gels (10% acrylamide), APB (5%) gel electrophoresis, and acidic PAGE.

### *In vitro* transcription

tRNA-Asp^(GUC)^ was prepared by run-off *in vitro* transcription using annealed DNA oligonucleotides as templates (**Supplementary Table S10**). Reactions containing 1x Transcription Buffer (ThermoFisher Scientific), 30 mM MgCl_2_, 5 mM DTT, 0.002 U/µL Pyrophosphatase (New England Biolabs), 6 mM rNTPs each, 40 ng/µL annealed template and 5 U/µL T7 RNA Polymerase (ThermoFisher Scientific) were performed at 37°C for 16 h followed by incubation with 0.1 U/µL TURBO™ DNase (ThermoFisher Scientific) at 37°C for 30 min. IVTs were ethanol precipitated before resuspension in IEX-A buffer (20 mM HEPES pH 7.5, 50 mM KCl) for anion-exchange chromatography. IVTs were loaded onto RESOURCE Q column (Cytiva), washed with 20% IEX-B buffer (20 mM HEPES pH 7.5, 2 M KCl) and eluted in a gradient of 20-35% buffer IEX-B. RNAs in the elution fractions were ethanol precipitated and resuspended in nuclease-free water. Fractions were pooled and separated by denaturing PAGE and RNAs were subsequently visualized by ultraviolet shadowing in TLC Silica gel 60 F_254_ plates (Merck Millipore). Relevant bands were excised and gel slices were crushed in gel extraction buffer (20 mM Tris-HCl, 250 mM sodium acetate, 1 mM EDTA, pH 5.0). Samples were flash-frozen at −80°C for 15 min followed by heating to 95°C for 5 min and incubation at room temperature for 3 h with mild rotation. Supernatants were recovered and the RNAs were ethanol precipitated and resuspended in nuclease-free water.

### Solid phase RNA synthesis and purification

RNA oligonucleotides were prepared by solid-phase synthesis using standard phosphoramidite chemistry with 2’-*O*-TOM-protecting groups on controlled pore glass solid support. Standard phosphoramidites, 5’-phosphate ON reagent and TBDMS-protected m^5^C phosphoramidite were purchased from ChemGenes and standard controlled glass solid supports were purchased from Sigma Aldrich. m^1^G phosphoramidite and 3’-phosphate CPG was prepared according to reported procedures ^69,70^. Automated solid phase synthesis was carried out on a K&A DNA/RNA synthesizer (H-6/H-8) on a 1 µmol scale. The following solutions were used: 0.25 m ethylthiotetrazole (ETT) in dry acetonitrile as activator, 3% dichloro acetic acid in 1,2-dichloroethane for detritylation, 50 mM iodine in water/pyridine 9/1 for oxidation, pyridine/acetic anhydride/THF 10/10/80 as Cap A solution and 16% NMI in THF as Cap B solution. RNA oligonucleotides were prepared on commercially available CPG solid supports using 70 mM solutions for standard and self-made phosphoramidites.

For unmodified sequences, deprotection was carried out with a mixture of aqueous ammonia and methylamine (1:1) at 37 °C for 4-6 h followed by removal of the 2’-silyl group by treatment with 500 µL of 1 M tetrabutylammonium fluoride in anhydrous THF at 37 °C for 14-16 h. After addition of 1 M Tris-HCl pH 8.0 (500 µL), the THF was removed in vacuum and the deprotected RNA was desalted by size exclusion chromatography using an ÄKTA start purification system with three HiTRAP Desalting columns (each 5 mL volume). The desalted oligonucleotides were eluted with water and a flow rate of 2 mL/min. The water was removed under reduced pressure and the samples were redissolved in 600 µL of nanopure water.

PAGE purification of the crude oligonucleotides was performed on 300×200×0.7 mm denaturing polyacrylamide gels (20% or 15% acrylamide/ bisacrylamide 19:1, 7 m urea) with 1x TBE buffer (89 mM Tris, 89 mM boric acid, 2 mM EDTA, pH 8.3). Gels were run at 35 W for 2 h 30 min. The oligonucleotides were visualized by UV shadowing on a TLC plate. After excision from the gel, the gel pieces were soaked in TEN buffer (10 mM Tris-HCl, 0.1 mM EDTA, 300 mM NaCl, pH 8.0) for oligonucleotide extraction and ethanol precipitated.

Crude as well as pure oligonucleotides were analyzed by anion exchange HPLC. A Dionex DNAPac PA200 column (2×250 mm) with buffer A (25 mM Tris-HCl, 6 M urea, pH 8.0) and buffer B (25 mM Tris, 0.5 M NaClO_4_, 6 M urea, pH 8.0) was used for this quality control. Samples were run with a gradient of 0-48% B over 12 column volumes with a flow rate of 0.5 mL/min and UV detection at 260 nm. The oven temperature was set to 60 °C. Pure oligonucleotides were additionally analyzed by HR-ESI mass spectrometry (micrOTOF-Q III, negative ion mode, direct injection) and the monoisotopic masses were obtained by deconvolution of the raw spectra. A summary of the oligonucleotides together with their respective measured and calculated molecular masses is provided in **Supplementary Table S11**.

### Ligation of tRNA fragments

tRNA^Tyr(GUA)^ was prepared by ligation of two fragments while tRNA^Asp(GUC)^ was accessed via a three-fragment splint ligation. For ligation of two fragments, 12.5 nmol of the 5′-phosphorylated fragment (R1107 or R1108) and 13.75 nmol of the 3’-OH fragment (R1106) were dissolved in water (87.5 µL) and annealed for 3 min at 95 °C. After cooling to ambient temperature, the solution was divided into two reaction vials (43.75 µL each) and 6.25 µL of 10 mM ATP, 6.25 µL of 10x T4 RNA ligase 1 buffer and 6.25 µL of T4 RNA ligase 1 were added to each vial. The reaction was incubated at 37 °C for 16 h. Afterwards, the solution was diluted 1:1 with high dye and the full-length tRNA product was purified by 10% PAGE. For tRNAs constructed from three fragments, first the 5’-fragment and the anticodon loop fragment were ligated in the presence of DNA splint 5 nmol of both the 5’-fragment R1112 and the anticodon loop fragment R1113 or R1128 were mixed with 4.2 nmol of DNA splint 1 (D3974), the volume was adjusted to 40 µL with water and the mixture was annealed for 3 min at 95 °C. After cooling to room temperature, 4.5 µL of 10x T4 RNA ligase 2 reaction buffer and 2.5 mL of T4 RNA ligase 2 were added. The reaction was incubated at 37 °C for 16 h, afterwards diluted 1:1 with high dye and purified by 15% PAGE. In the second step, 1.8 nmol of the first ligation product was combined with 1.8 nmol of the 3’-fragment R1114 and 1.5 nmol of DNA splint 2 (D3975) in a volume of 16 µL water. The mixture was annealed for 3 min at 95 °C, cooled to room temperature, 2 µL of 10x T4 RNA ligase 2 reaction and 2 µL of T4 RNA ligase 2 were added and the mixture was incubated at 37 °C for 16 h. The full-length tRNA was obtained by 10% PAGE purification. For dephosphorylation, 1 nmol of full-length tRNA was dissolved in 34 µL water, 4 µL of 10x rCut Smart buffer and 2 µL of CIAP were added. The mixture was incubated for 30 min at 37 °C. The dephosphorylated tRNA was extracted with a phenol/chloroform/ isoamylalcohol mixture, precipitated with 70% ethanol and the dried pellet was redissolved in water.

### Recombinant expression and purification of TGT

Expression of recombinant TGT was performed by co-expression of the QTRT1 and QTRT2 subunits in *E. coli* BL21(DE3) as described previously ^12^. Protein expression was performed by autoinduction in ZYM-5052 medium for 3 h at 37 °C followed by 60-65 h at 16 °C. Recombinant TGT was purified via a His_6_ tag on QTRT1. Bacterial pellets were resuspended in lysis buffer (50 mM HEPES pH 7.5, 100 mM NaCl, 5 mM imidazole) supplemented with cOmplete EDTA-free Protease Inhibitor Cocktail (Roche) and disrupted in a microfluidizer unit (EmulsiFlex-C3, Avestin.Inc) by five passages at 10,000 psi. Crude lysates were centrifuged at 20,000 xg for 1 h at 4 °C and the supernatant was filtered at 0.45 µm for subsequent loading onto HiTrap TALON Crude columns (Cytiva). Columns were equilibrated in loading buffer (50 mM HEPES pH 7.5, 100 mM NaCl, 5 mM imidazole) and thoroughly washed with LiCl wash buffer (50 mM HEPES pH 7.5, 100 mM NaCl, 2 M LiCl) after sample binding. TGT was eluted in 25% elution buffer (50 mM HEPES pH 7.5, 100 mM NaCl, 500 mM imidazole). Elution fractions containing TGT were pooled and concentrated with 10 kDa molecular weight cut-off (MWCO) Amicon Ultra Centrifugal Filters (Merck Millipore). The complex was further purified by size-exclusion chromatography (SEC) using HiLoad 26/600 Superdex 200 pg column (Cytiva) equilibrated in SEC buffer (20 mM HEPES pH 7.5, 100 mM NaCl) and fractions containing TGT were concentrated as above. The protein was quantified with the Pierce BCA Protein Assay kit (ThermoFisher Scientific) and purity was assessed by SDS-PAGE followed by staining with Coomassie R-250.

### *In vitro* queuosinylation

Reactions containing 2 µg recombinant TGT, 1 µg tRNA and 1 mM queuine base were incubated at 37 °C for 1 h in Q-incorporation buffer (100 mM HEPES pH 7.5, 20 mM MgCl_2_, 5 mM DTT). RNAs were extracted with TRI Reagent LS (Sigma-Aldrich) and Q incorporation was confirmed by 0.5% ABP gel electrophoresis followed by northern blotting or SYBR Gold staining as appropriate. The Q incorporation rate was determined by calculating the ratio of Q-containing tRNA to total tRNA (Q- and G-containing tRNA) in each sample. Images were analyzed and quantified in ImageStudio Lite software (LICORbio).

### Recombinant expression and purification of ANG and its derivatives

Expression of recombinant ANG and the corresponding mutants from plasmids containing a codon-optimized sequence encoding ANG lacking the N-terminal signal peptide was performed in BL21-CodonPlus (DE3)-RIL *E. coli* in TB medium at 37°C for 3 h with 1 mM IPTG. Proteins were recovered from inclusion bodies and refolded as described previously ^71^. Bacterial pellets were resuspended in lysis buffer (50 mM Tris-HCl pH 8.0, 2 mM EDTA) and disrupted in a microfluidizer unit (EmulsiFlex-C3, Avestin.Inc) by five passages at 10,000 psi. Lysates were centrifuged at 49,000 xg for 20 min at 4°C and the pellets containing insoluble protein were washed in lysis buffer supplemented with 5% (v/v) Triton X-100 to remove membrane proteins and lipids. Pellets were sonicated in a Bandelin Electronic Sonopuls HD2070 device to homogenize the suspension and centrifuged as described above. The resulting pellets were washed in lysis buffer to remove residual detergent followed by another round of sonication and centrifugation as described above. The final pellets were resuspended in solubilization buffer (7 M guanidine-HCl, 0.15 M reduced glutathione, 2 mM EDTA pH 8.0) and gently stirred for 3 h to solubilize inclusion bodies. Samples were subsequently filtered through a 0.45 µm syringe filter to remove remaining insoluble material. Refolding was performed by dropwise dilution of filtered samples in 500 mM arginine-HCl pH 8.0 supplemented with 0.6 mM oxidized glutathione under gentle stirring for 30 min. Refolded samples were diluted 4-fold in water to reduce salinity and loaded onto HiTrap SP Sepharose FF columns (Cytiva) for ion-exchange chromatography. Columns were equilibrated in buffer A (25 mM Tris-HCl pH 8.0, 0.2 M NaCl) and proteins were eluted in a gradient of 0-50% buffer B (25 mM Tris-HCl pH 8.0, 2 M NaCl). Elution fractions containing the target protein were pooled and concentrated using 3 kDa MWCO Amicon Ultra Centrifugal Filters (Merck Millipore) before further purification by size-exclusion chromatography (SEC) using a HiLoad Superdex 16/600 75 pg column (Cytiva) in SEC buffer (20 mM Tris-HCl pH 8.0, 120 mM NaCl). The protein was quantified with the Pierce BCA Protein Assay kit (ThermoFisher Scientific) and purity was assessed by SDS-PAGE in 12% Bis-Tris gels followed by staining with Coomassie R-250.

### *In vitro* tRNA cleavage assays

Reactions were carried out by mixing 0.5 µg unmodified or Q-modified tRNA-Asp^(GUC)^ *in vitro* transcript and 0.625 µg/mL recombinant ANG (or its mutant versions) in cleavage buffer (30 mM HEPES pH 6.8, 30 mM NaCl, 5 mM MgCl_2_, 0.1 mg/mL BSA) ^72^. Alternatively, 1 µg small RNAs and 1.25 µg/mL ANG_Q117G_ or ANG_H114A/Q117G_ were used. Samples were incubated at 37°C for 30 min and reactions were quenched by adding equal volume of RNA Loading Dye (2×). Cleavage products were separated on 12% denaturing (7 M urea) polyacrylamide TBE gels, and RNAs were detected by SYBR Gold staining and/or northern blotting.

## DATA AVAILABILITY

The bisulfite sequencing data generated during this study have been deposited in Gene Expression Omnibus database [http://www.ncbi.nlm.nih.gov/geo/] under the following accession code: GSE334738.

## ACKNOWLEDGEMENTS

This work was funded by the Deutsche Forschungsgemeinschaft (DFG, German Research Foundation), project number 469281184, SFB1565 to RF (P09), KEB (P12), CH (P18), MTB (P18).

The authors gratefully acknowledge the computing time granted by the Resource Allocation Board and provided on the supercomputer Emmy/Grete at NHR-Nord@Göttingen as part of the NHR infrastructure. The calculations for this research were conducted with computing resources under the project scc_ukbm_ bohnsack.

## SUPPLEMENTARY INFORMATION

**Supplementary Figure S1.**
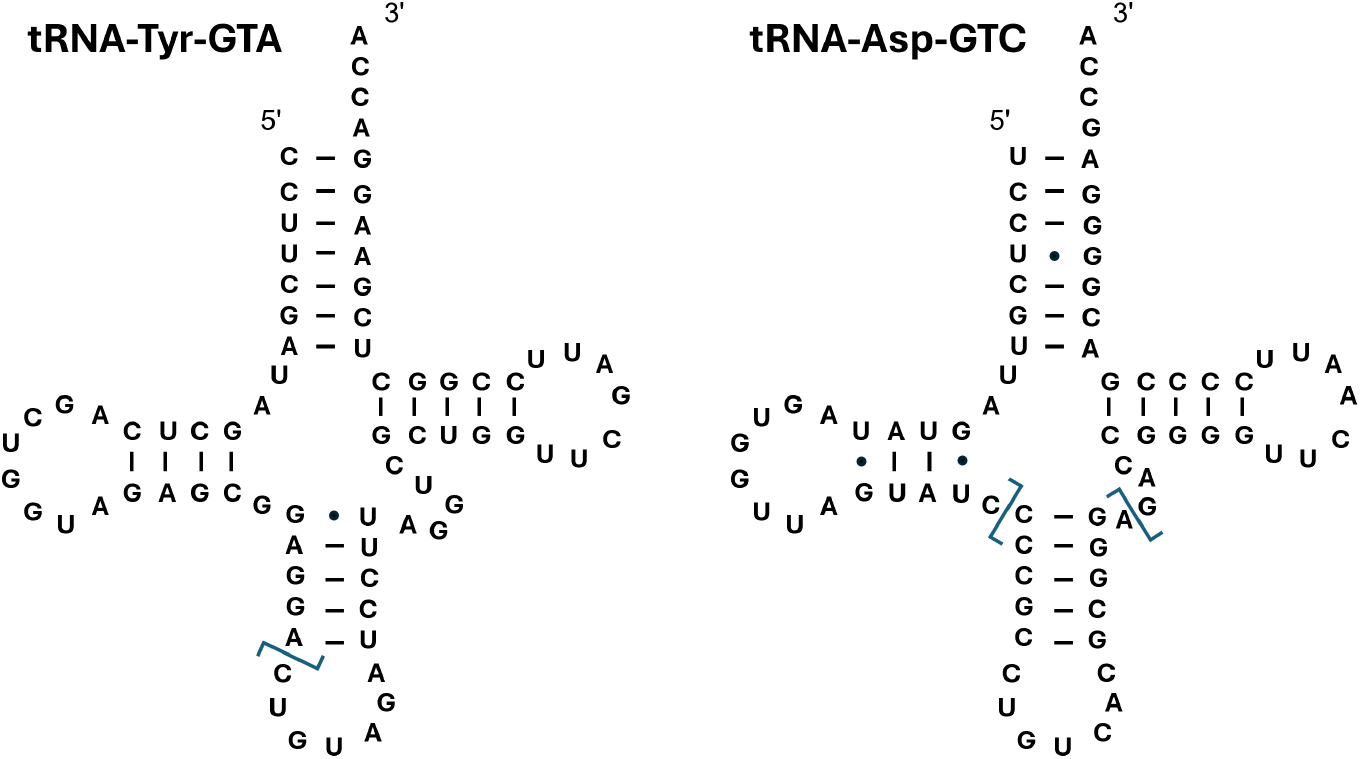
Secondary structures of tRNA^Tyr(GUA)^ and tRNA^Asp(GUC)^ with the ligation sites indicated by blue lines.

**Supplementary Figure S2.**
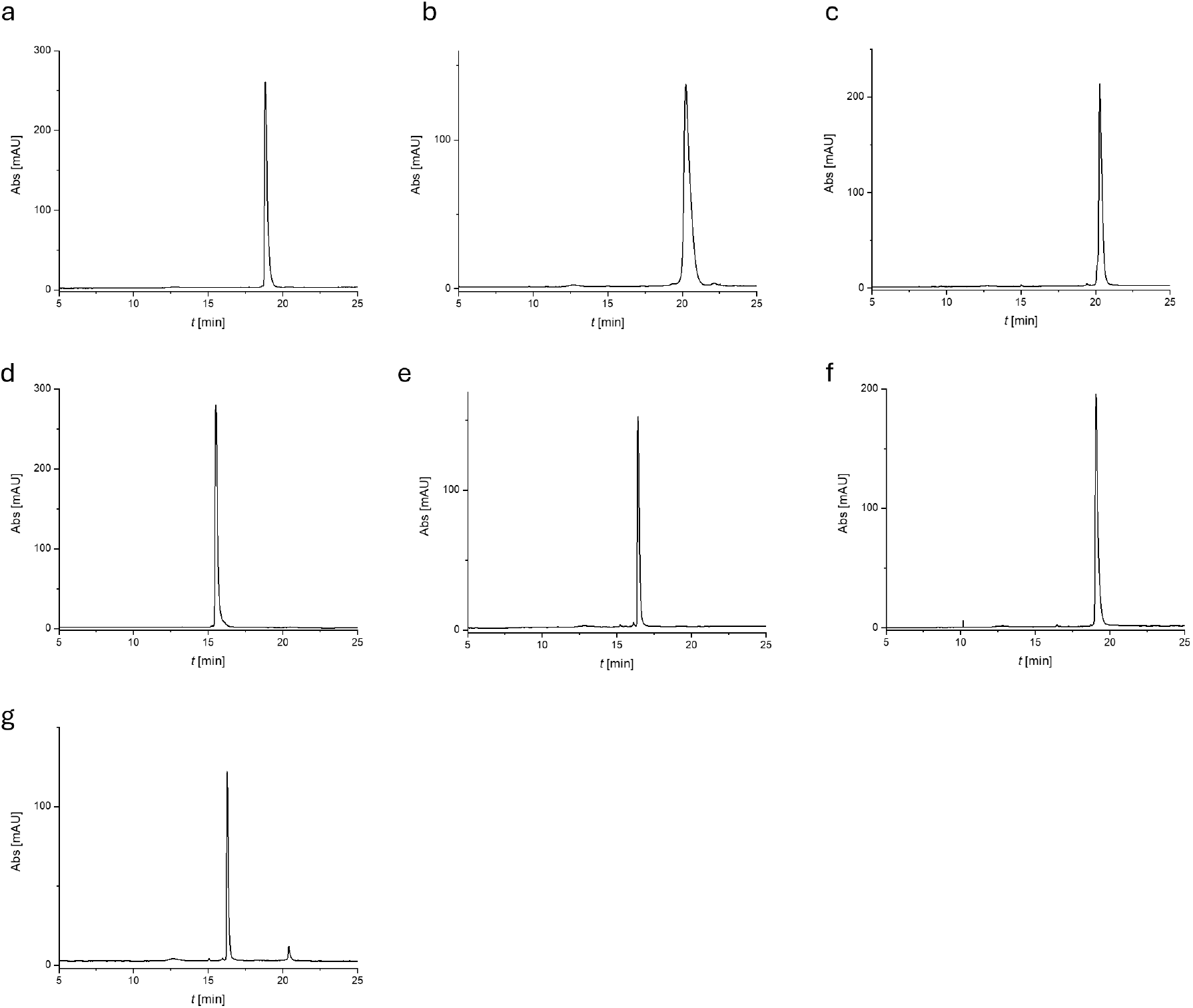
HPLC chromatograms for purified synthetic RNA oligonucleotides: a) R1106, b) R1107, c) 1108, d) R1112, e) R1113, f) R1114 and g) R1128 (see table EV1 for RNA oligo details; below) Anion exchange HPLC, Dionex, DNAPac PA200, 2×250mm, 60 °C.

## SUPPLEMENTARY TABLES

**Supplementary Table S1.**
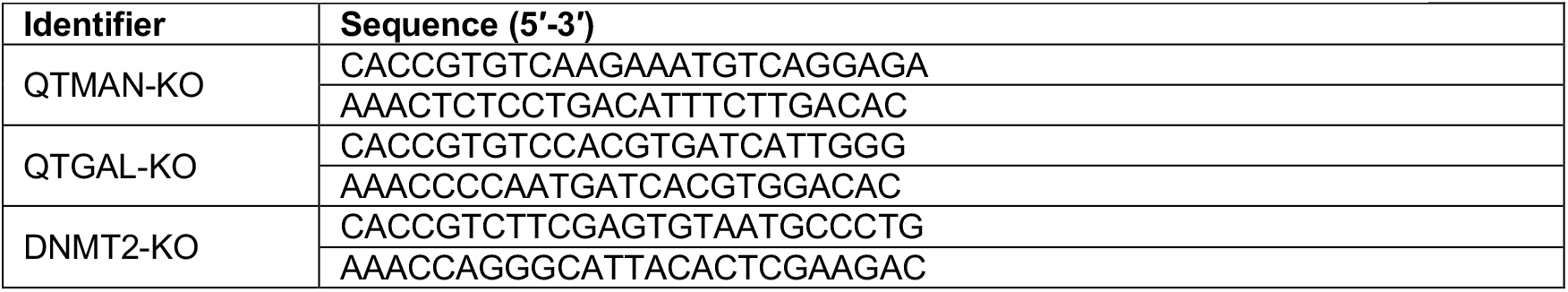
Oligonucleotides used for CRISPR-Cas9 sgRNA assembly.

**Supplementary Table S2.**
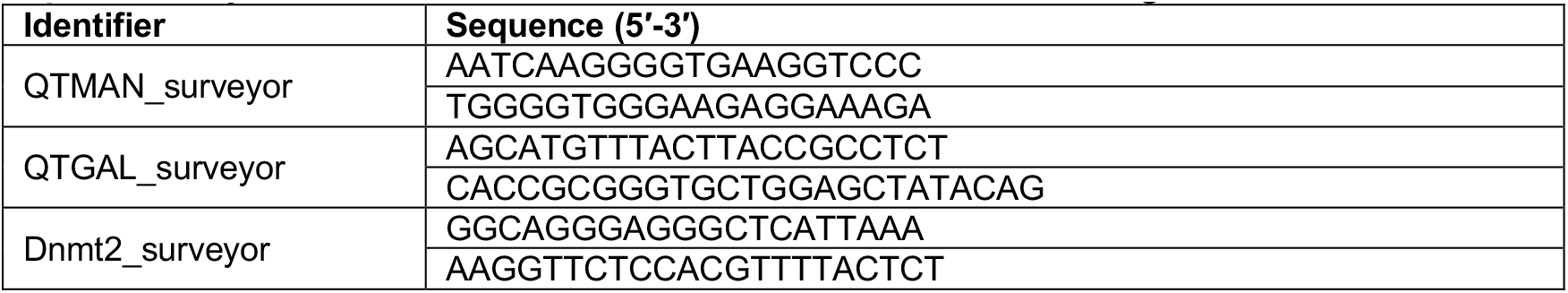
DNA oligonucleotides used for CRISPR-Cas9 genome amplification.

**Supplementary Table S3.**
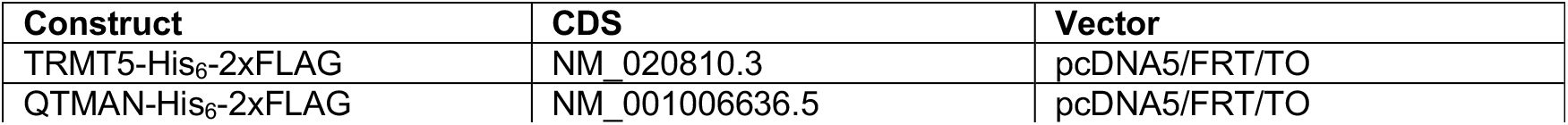
Mammalian expression plasmids used in this study.

| Construct | CDS | Vector |
| --- | --- | --- |
| TRMT5-His<sub>6</sub>-2xFLAG | NM_020810.3 | pcDNA5/FRT/TO |
| QTMAN-His<sub>6</sub>-2xFLAG | NM_001006636.5 | pcDNA5/FRT/TO |

**Supplementary Table S4.**
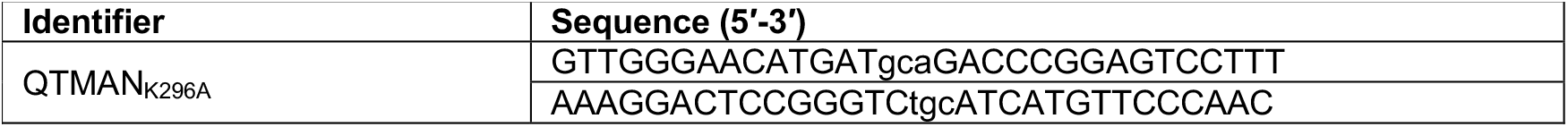
DNA oligonucleotides used for site-directed mutagenesis.

**Supplementary Table S5.** Primary antibodies used in this study.

| Target | Supplier | Dilution |
| --- | --- | --- |
| α-Tubulin | Sigma-Aldrich (T6199) | 1:5,000 |
| 6*His-tag | Proteintech (66005-1-Ig) | 1:5,000 |
| DNMT2 | Proteintech (19221-1-AP) | 1:500 |
| FLAG | Sigma-Aldrich (F3165) | 1:10,000 |
| GAPDH | Proteintech (10494-1-AP) | 1:25,000 |

**Supplementary Table S6.** Secondary antibodies used in this study.

| Secondary antibody | Supplier | Dilution |
| --- | --- | --- |
| IRDye680LT Donkey anti-Mouse | LICORbio | 1:10,000 |
| IRDye800CW Donkey anti-Mouse | LICORbio | 1:10,000 |
| IRDye680RD Donkey anti-Rabbit | LICORbio | 1:10,000 |
| IRDye800CW Donkey anti-Rabbit | LICORbio | 1:10,000 |

**Supplementary Table S7.**
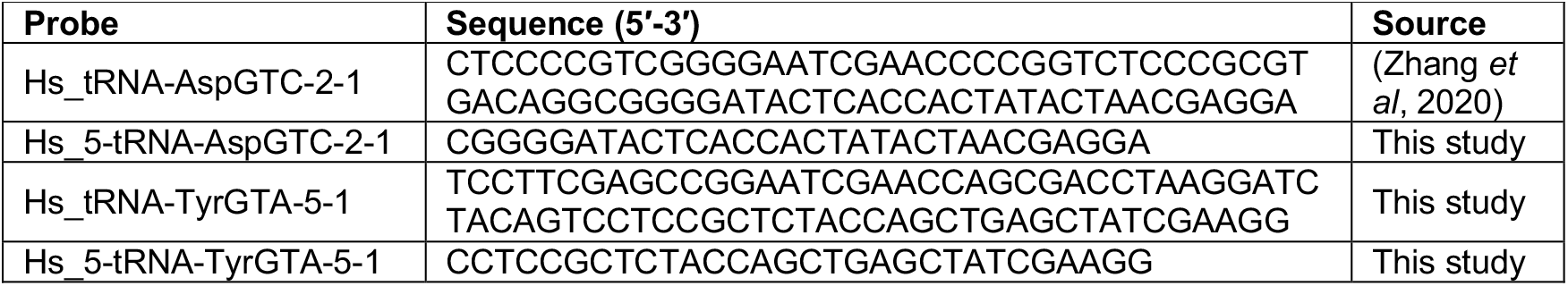
Northern blotting probes used in this study.

**Supplementary Table S8.**
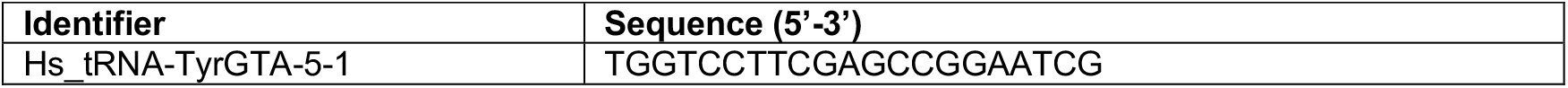
DNA oligonucleotides used for primer extension.

**Supplementary Table S9.**
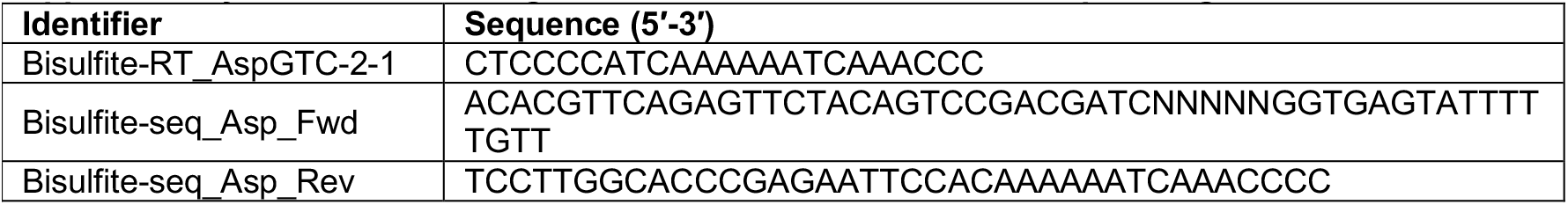
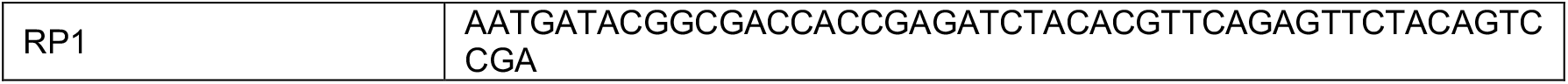
DNA oligonucleotides used for bisulfite sequencing.

**Supplementary Table S10.**
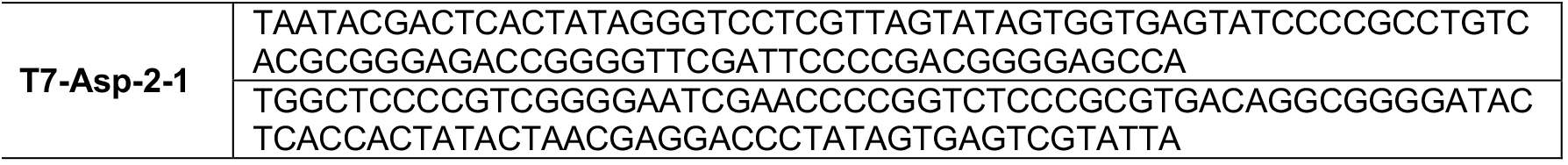
DNA oligonucleotides used for *in vitro* transcription.

**Supplementary Table S11.**
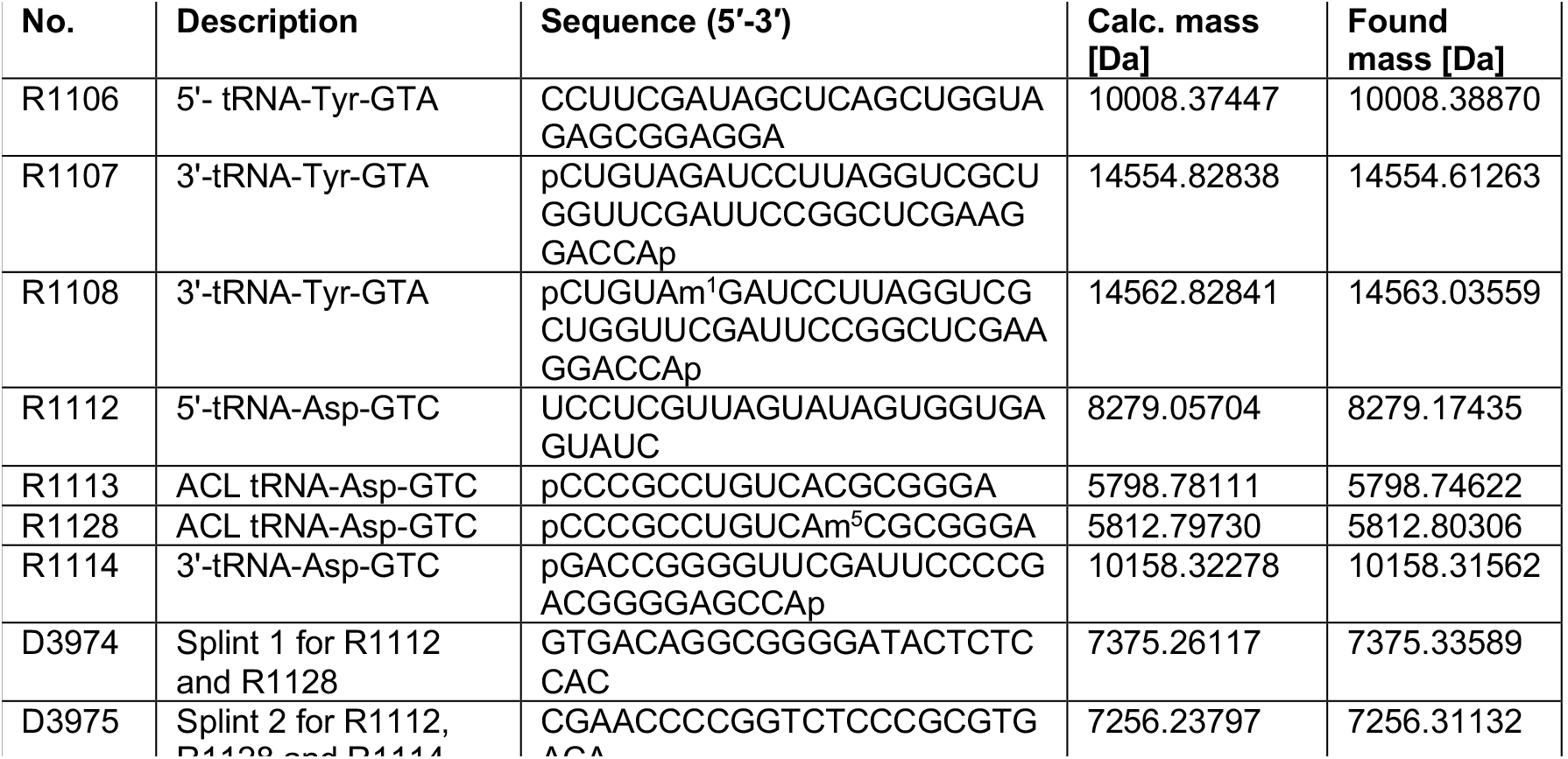
Sequences of RNA oligonucleotides prepared by solid phase synthesis and of the DNA splints with calculated and measured HR-ESI-MS masses.

## SUPPLEMENTARY METHOD

### Bisulfite data analysis

For samples prepared from total RNA, quality control analysis of data was performed using FASTQC (v.0.11.5) and paired-end FASTQ files were assembled using PEAR (v0.9.6) with the following parameters: -p 0.05 -n 4 -v 6. Due to library multiplexing during sequencing, R2 was used as the forward-oriented read to reconstruct the expected amplicon sequence. Assembled reads were then filtered based on the expected PCR product length using custom Bash scripts implemented with awk (v5.0.1). The 5-nucleotide random barcode (NNNNN) and additional five nucleotides at the 5’ end were trimmed using Cutadapt (v4.9) with the settings -u 10 -j 4. The resulting FASTQ files were again assessed for base composition with FASTQC and quality metrics were summarized using MultiQC (v1.13.dev0).

For input and IP eluate samples, quality control of the raw sequencing data was performed using FastQC (v.0.11.9), and the resulting quality metrics were summarized using MultiQC. Paired-end reads were assembled using PEAR (v0.9.6) with the parameters ‘-p 0.05 -n 4 -v 6’, followed by an additional FastQC and MultiQ (v1.12) assessment of the assembled reads. Adapter and quality trimming were subsequently performed using Flexbar (v3.5.0) with right-end trimming, a minimum read length of 15 nucleotides, a quality threshold of 13, and the adapter sequence “TGGAATTCTCGGGTGCCAAGG”. Reads were then filtered to retain sequences within the expected product-length range using a custom Bash script based on awk, after which sequence quality was reassessed using FastQC and MultiQC. Barcode removal was performed using pyBarcodeFilter.py (v3.1)^1^ with a predefined barcode NNNNNGGT, and the resulting barcode-filtered reads were subjected to further quality-control analysis.

